# P1 bacteriophage-enabled delivery of CRISPR-Cas9 antimicrobial activity against *Shigella flexneri*

**DOI:** 10.1101/2022.09.02.506314

**Authors:** Yang W. Huan, Vincenzo Torraca, Russell Brown, Jidapha Fa-arun, Sydney L. Miles, Diego A. Oyarzún, Serge Mostowy, Baojun Wang

## Abstract

The discovery of clustered, regularly interspaced, short palindromic repeats (CRISPR) and the Cas9 RNA-guided nuclease provides unprecedented opportunities to selectively kill specific populations or species of bacteria. However, the use of CRISPR-Cas9 to clear bacterial infections *in vivo* is hampered by the inefficient delivery of *cas9* genetic constructs into bacterial cells. Here, we use a broad-host-range P1-derived phagemid to deliver the CRISPR-Cas9 chromosomal-targeting system into *Escherichia coli* and the dysentery-causing *Shigella flexneri* to achieve DNA sequence-specific killing of targeted bacterial cells. We show that genetic modification of the helper P1 phage DNA packaging site (*pac*) significantly enhances the purity of packaged phagemid and improves the Cas9-mediated killing of *S. flexneri* cells. We further demonstrate that P1 phage particles can deliver chromosomal-targeting *cas9* phagemids into *S. flexneri in vivo* using a zebrafish larvae infection model, where it significantly reduces the bacterial load and promotes host survival. Our study highlights the potential of combining a P1 bacteriophage-based delivery with the CRISPR chromosomal-targeting system to achieve DNA sequence-specific cell lethality and efficient clearance of bacterial infection.

## INTRODUCTION

CRISPR (clustered regularly interspaced short palindromic repeats) in combination with the Cas (CRISPR-associated) endonuclease enzyme(s) constitutes the immune system of prokaryotes, serving to protect bacteria against invading nucleic acids^1,2^. Cas9 is an RNA-guided endonuclease that introduces double strand cleavage at its target DNA sequence, while the specificity is determined by the spacer sequence of CRISPR RNA (crRNA), which is complementary to the target DNA sequence^3,4^. Previous studies demonstrated that the CRISPR-Cas9 system can be programmed to target antibiotic-resistance gene(s) or species-specific chromosomal gene(s) of bacteria. Such targeting can re-sensitise bacterial populations to antibiotic treatment or, in the latter case, induce cell lethality via SOS-mediated responses against double-stranded DNA damage in the bacterial chromosome. Cell lethality via DNA sequence-specific targeting has been described for various clinically relevant bacterial pathogens, such as antimicrobial-resistant strains of *Escherichia coli*^5,6^, *Staphylococcus aureus*^7^ and *Salmonella enterica spp*^8^. Despite this success, the use of CRISPR-Cas9 as an antimicrobial system is hampered by the low transformation efficiency of target bacteria, especially during infections *in vivo*^6^.

Bacteriophages, viruses that predate on bacterial cells, have been applied in treating bacterial infections for over a century^9,10^. New strains of lytic bacteriophages recovered from environmental and biological samples are routinely used in studies to kill clinically relevant and/or multidrug-resistant strains of bacteria, including *Clostridium difficile*^11^, *Shigella flexneri*^12^, *Pseudomonas aeruginosa*^13^ and methicillin-resistant *Staphylococcus aureus* (MRSA)^14^. While the direct use of lytic bacteriophage cocktails is not widely accepted as a reliable alternative to antibiotics in the clinical field, properties of bacteriophages such as specific host range, high transduction efficiency, stability of bacteriophage particles and the ability of certain bacteriophages to lysogenise into host cells make them suitable as an efficient delivery tool for genetic constructs^15,16^. Citorik et al demonstrated M13 phagemid-based delivery of the Cas9 system into carbapenem-resistant *E. coli* to target its antibiotic resistance determinants *bla*NDM-1 or *bla*SHV-18, which allowed re-sensitisation of the bacteria towards antibiotic treatment *in vitro*^17^. In addition, the Cas9 endonuclease was reprogrammed to target a chromosomally encoded virulence factor (*eae*) of enterohemorrhagic *E. coli*, which allowed DNA sequence-specific killing of the bacteria in a *Galleria mellonella* infection model^17^. Bikard et al., demonstrated the use of Φ NM1 bacteriophage to deliver phagemid-encoded CRISPR Cas9 antimicrobial systems into *S. aureus*^5^. In this case, reprogramming of the Cas9 endonuclease to target the methicillin resistance gene, *mecA*, was introduced to specifically kill the methicillin-resistant strain USA300Φ, but not the RNΦ strain, *in vitro*.^5^ The sequence-specific killing effect of Cas9 was expanded to target the chromosomal *aph* gene of RNKΦ strain, which selectively reduced 40% of the RNKΦ cells in vivo using a mouse skin colonisation model ^5^. Recently, the M13 phagemid-based delivery of the Cas9 system allowed the selective killing of a F^+^ *E. coli* strain in a murine gut colonisation model^18^. In this case, the authors demonstrated a selective reduction of a *gfp*^+^ F^+^ *E. coli* strain (approximately 1-3 log), using a *gfp*-targeting M13 *cas9* phagemid^18^. Taken together, these studies have suggested that bacteriophages allow efficient delivery of DNA sequence-specific Cas9 antimicrobials into bacteria *in vivo*, and strongly support the approach as an alternative therapeutic option to treat bacterial infections.

Shigellosis is an acute intestinal infection caused by *Shigella* spp. Worldwide, it was estimated that *Shigella* caused 80–165 million cases of disease and 600,000 deaths annually^19^. *S. flexneri* is most frequently recorded in developing countries, with a disproportionately high mortality rate in children^20,21,22^. Clinical isolates of *S. flexneri* are often drug resistant, and it was estimated that half of all contemporary strains of *Shigella* spp are multidrug resistant^23^. This calls for alternative therapeutic options, such as the use of bacteriophage in treating *Shigella* infection. In this study, we demonstrate the use of a broad-host-range transducing P1 phagemid system to deliver chromosomal-targeting *cas9* genetic constructs into *E. coli* and the dysentery-causing *S. flexneri* to achieve sequence-specific lethality of targeted bacterial cells. To reduce the packaging of the resident P1 bacteriophage genome, we performed genetic modification of the *pac* site, the recognition site for the pacase enzyme, to bias the packaging of *cas9* phagemid into P1 phage particles. We show that improved titre of P1 transducing units in the lysates prepared from *pacA*\*::*npt* EMG16 cells significantly enhanced the Cas9-mediated lethality effect on *S. flexneri*. Moreover, we show that the *cas9* chromosomal-targeting phagemid is efficient in inducing sequence-specific lethality of *S. flexneri in vivo*, and significantly improved the survival of infected zebrafish larvae. Overall, these results highlight the potential of P1-based phagemid delivery of the chromosomal-targeting CRISPR-Cas9 system as a powerful tool to target clinically relevant and antibiotic-resistant Gram-negative *Enterobacteriaceae* pathogens.

## RESULTS

### Construction of P1 BBa_J72114 phagemid with high transduction efficiency in *E. coli* strains

To deliver and stably express foreign genetic cassette(s) in *Enterobacteriaceae*, the BBa_J72114 P1 phagemid constructed for this study contains all necessary P1-based elements for packaging of phagemids into P1 bacteriophage particles (refer to **Supplementary Figure 1** for details), as well as a selectable chloramphenicol-resistance marker and p15A or pBBR1 origin of replication for phagemid maintenance (**Figure 1a**).

**Figure 1:**
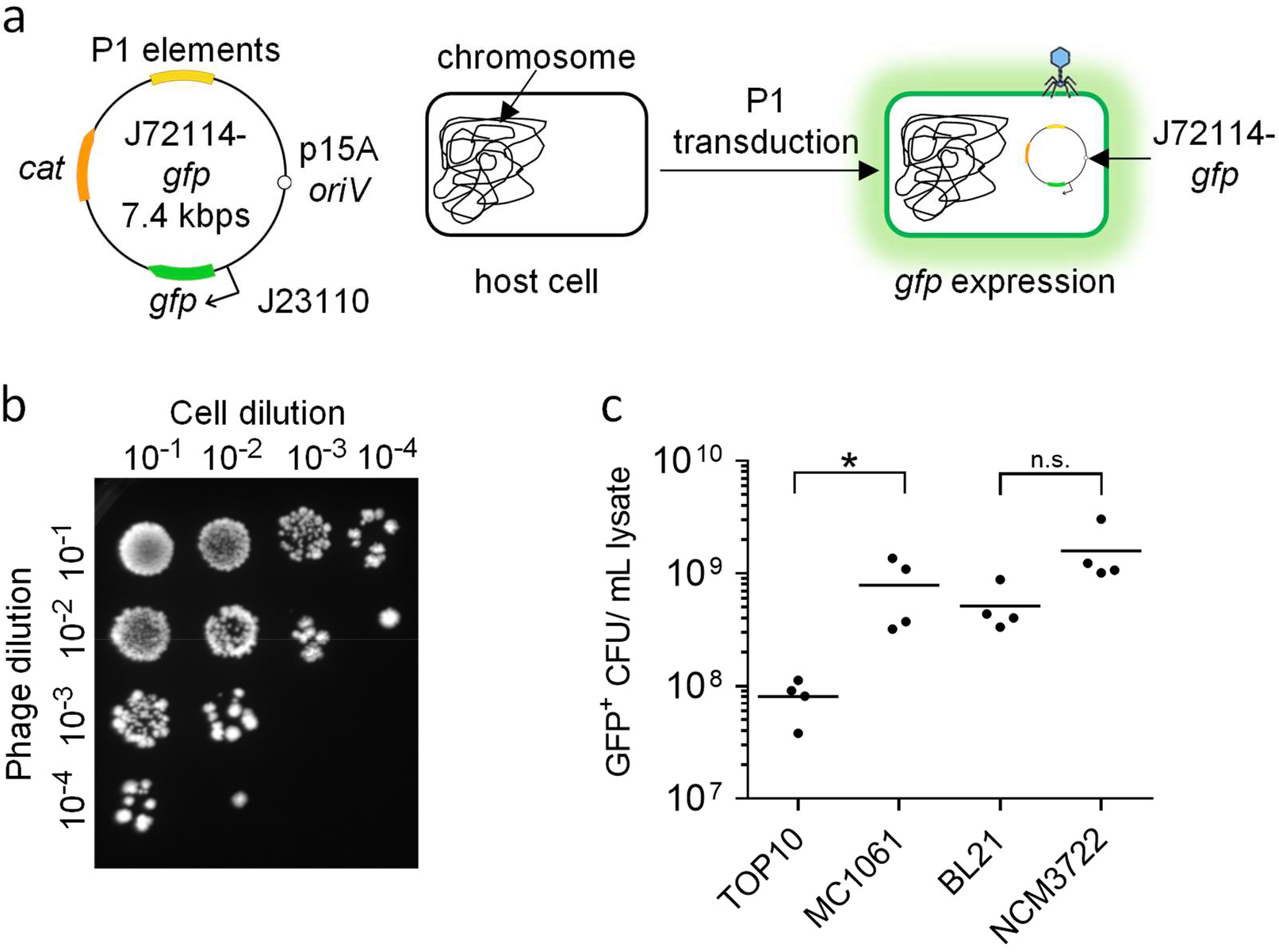
P1 J72114 phagemid as a delivery tool for transduction of foreign genetic cassette into Gram-negative *Enterobacteriaceae*. (**a**) Schematic diagram of P1-J72114 phagemid, characterised by constitutive expression of *gfp* placed under the BBa_J23115 promoter. The phagemid contains the chloramphenicol acetyltransferase gene (*cat*), which confers a chloramphenicol-resistant phenotype to transduced or transformed cells. Transduced *E. coli* cells will retain the J72114-*gfp* phagemid, giving constitutive *gfp* expression. (**b**) Representative image showing the presence of GFP-positive *E. coli* NCM3722 cells after transduction with serially diluted phagemid lysates delivering the *gfp* expression cassette. Serial dilutions of recovered cells (10^1^, 10^2^, 10^3^, 10^4^) were made and spotted onto LB agar supplemented with chloramphenicol. (**c**) Quantification of transducing units (phage particles containing phagemid sequences) in various lab strains of *E. coli*. Each data point represents a biological replicate and is the average of 4 technical repeats. Horizontal lines represent the group mean. The p-values were determined by one-way ANOVA, and significance was defined as p < 0.05, shown as *, while n.s. represents no significant difference(s).

Due to the non-replicative nature of the P1 phagemid, an excess of phagemid is required to significantly impact the target bacterial population. To quantify the P1 transducing units’ titre of lysates prepared, the original phagemid was modified to include a constitutive *gfp* expression cassette (J72114-*gfp*, **Figure 1a**). Various lab strains of *E. coli* were transduced with P1 phage lysates and recovered cells were selected for chloramphenicol resistance and GFP fluorescence (**Figure 1b**). The P1 lysates transduced all 3 sub-strains of *E. coli* K-12 and*E. coli* BL21 within a range of 7 × 10^8^ to 7 × 10^9^ transducing units per millilitre of lysate used (**Figure 1c**). Overall, our protocol generates sufficient phagemid titre for the delivery and stable expression of genetic constructs into *E. coli*.

### The efficacy of Cas9-induced cell lethality of *E. coli* MC1061::*npt*

We constructed J72114-*cas9* phagemid to achieve sequence-specific DNA cleavage on target enterobacterial cells. The complete Cas9 system was derived from p*Cas9* plasmid with constitutive expression of the *cas9, trans*-activating RNA (*tracRNA*) and CRISPR RNA (*crRNA*) (**Figure 2a**). The specificity and efficacy of Cas9 endonuclease-mediated lethality was first evaluated on *E. coli* K12 strain MC1061::*npt*, which has a single copy of chromosomally integrated *npt* gene conferring kanamycin resistance. The *npt*-targeting spacer sequence was cloned into *cas9* phagemid using a BsaI cloning system (refer to **Supplementary Table 3** for DNA sequences). Since *E. coli* MC1061::*npt* is not a recombination deficient mutant, chromosomal DNA double-strand breaks (DSBs) caused by Cas9 endonuclease(s) in the presence of chromosomal-targeting spacer sequence(s) would induce an SOS-mediated response, leading to DNA repair, cell cycle arrest and/or apoptosis-like cell death^33^ (**Figure 2a**). This would lead to a reduced CFU recovery after treatment with *cas9*-*npt* phagemid, as compared to that of *cas9* phagemid without the *npt*-targeting spacer sequence, as well as mock-infected cells.

**Figure 2:**
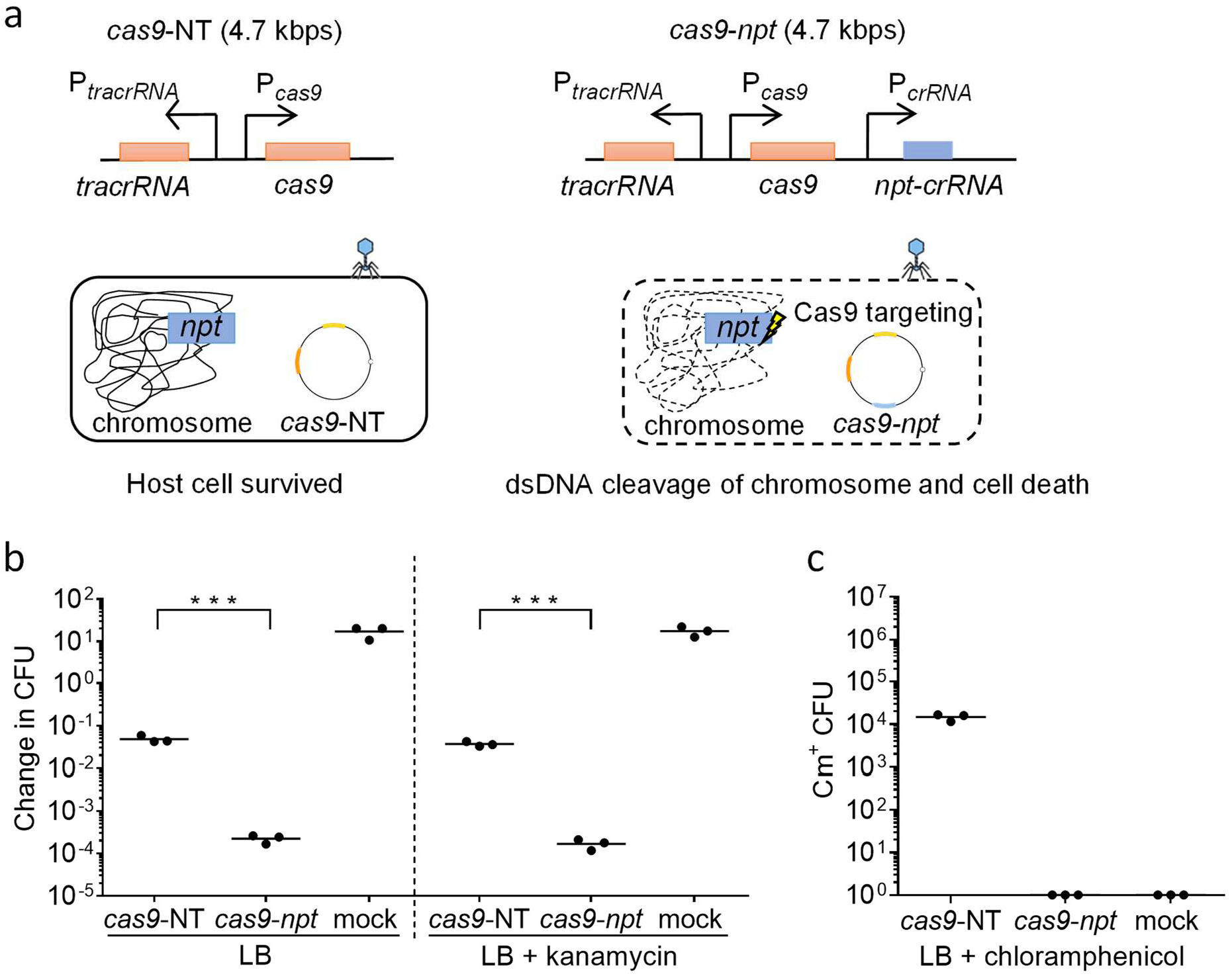
Spacer sequence mediated lethality of *E. coli* MC1061::*npt* cells using *npt*-targeting *cas9* phagemid. (**a**) Schematic diagrams showing the *cas9* genetic construct, with or without *npt*-targeting crRNA (*cas9*-*npt* and *cas9*-NT respectively) assembled onto the P1 J72114 phagemid. The presence of *npt-*targeting *crRNA* would target the Cas9 endonuclease chromosome of *E. coli* MC1061::*npt* cells, causing dsDNA cleavage of the chromosome and cell death. (**b**) Serial dilutions of transduced *E. coli* MC1061::*npt* were plated onto plain LB agar or LB agar supplemented with kanamycin. Data were plotted as change(s) in CFU as compared to input CFU (approximately 10^7^ cells per reaction) used for infection. (**c**) Quantification of chloramphenicol-resistant CFUs recovered, after treatment with *cas9*-NT or *cas9*-*npt* phagemid lysates. Each data point represents a biological replicate and is the average of 4 technical repeats. MOI of 5 P1 transducing units to 1 bacterial cell was used for all infections. Horizontal bars represent the group mean. The p-values (between non-targeting and targeting phagemid treatments) were determined using a two-tailed unpaired t-test with significance defined by p < 0.05. p < 0.0005 between targeting phagemid and non-targeting phagemid treatments are shown as ***.

The killing effect mediated by the presence of *npt*-targeting spacer sequence (ΔCFUNT-*npt*) was ∼100-fold (p < 0.0005) higher than that elicited by non-targeting *cas9*-NT phagemid (**Figure 2b**). The CFU recovered after treatment with *cas9*-*npt* phagemid was not significantly different in the presence or absence of kanamycin (CFU recovered on plain LB: 65.01 ± 4.97; CFU recovered on LB+kanamycin: 57.14 ± 4.08, p > 0.05), suggesting that the reduction in CFU is due to cell lethality caused by Cas9 chromosomal-targeting activity and not to the loss of the *npt* gene and/or its gene function (**Figure 2b**). These results showed that non-targeting *cas9*-NT phagemid treatment caused ∼25-fold and ∼250-fold reduction in recovered CFU as compared to an input of ∼10^7^ CFU and ∼10^8^ CFU recovered from the mock infection, respectively (**Figure 2b**). The non-Cas9 spacer sequence mediated killing effect may be attributed to the general cytotoxic effect of lysates. We did not recover any chloramphenicol-resistant cells after treatment with *cas9-npt* phagemid, suggesting that the presence of both *npt* gene and *npt*-targeting spacer sequence of the *cas9* phagemid would always lead to cell death (**Figure 2c**). There was no significant difference between CFU recovered after *cas9*-*npt* or *cas9*-NT phagemid treatment of *E. coli* K12 MC1061 cells without the chromosomal *npt* gene (**Supplementary Figure 2**). This verified that the specificity of Cas9-mediated lethality depends on the presence of both the *npt*-targeting spacer sequence and the chromosomal *npt* gene sequence.

Overall these results show that the *cas9* phagemid with chromosomal-targeting spacer sequence is unstable or conditionally lethal when introduced into target bacterial cells.

### Genetic modification of *pac* site of P1 genome to improve P1 phagemid purity

Previous results showed that P1 phagemid lysates confer general cytotoxicity on *E. coli* K12 MC1061 cells, irrespective of the presence or absence of the Cas9 spacer sequence. The non-spacer sequence mediated lethality effect (termed general cytotoxicity of lysate) may be attributed to the presence of wildtype P1 phage in the lysates, which are capable of undergoing lytic stage replication hence the killing of transduced cells. This prompted us to genetically remove the DNA packaging site, *pac*, on the resident wildtype P1 phage to bias the packaging of cas9 phagemid and reduce wildtype P1 phage titre. The 161 bp *pac* sites lie within *pacA* gene sequence and contain seven hexameric repeats (“TGATCA/G”) with “GATC” Dam methylation site^34,35^. Previous studies proposed that the hemimethylated *pac* site would be recognised and bound by the pacase enzyme, while further methylation would promote cleavage of the *pac* site by the bound pacase^34,35,36^. We hypothesised that disruption of these hexamer repeat motifs would reduce the packaging of the wildtype P1 genome. The *E. coli* P1 lysogen EMG16 strain with a modified *pacA* gene sequence (termed *pacA\**::*npt)* was created by introducing synonymous mutations into the hexameric repeats of the *pac* site via lambda-red recombineering (**Figure 3a**). Phage lysates of *cas9* phagemid without chromosomal-targeting spacer sequences were prepared from wildtype and *pacA\**::*npt* EMG16 cells. Quantification of plaque-forming units and transducing units suggested that lysates prepared from the *pacA\**::*npt* mutant contained approximately 9-fold lower wildtype P1 phage titre compared to that of lysates prepared from wildtype EMG16 cells (p < 0.0005, **Figure 3b**). There was no significant difference in the phagemid titres of lysates prepared from both wildtype and *pacA\**::*npt* EMG16 cells, indicating that the *pacA* genetic modification had negligible effects on the packaging of phagemid into transducing units (p > 0.05, **Figure 3b**).

**Figure 3:**
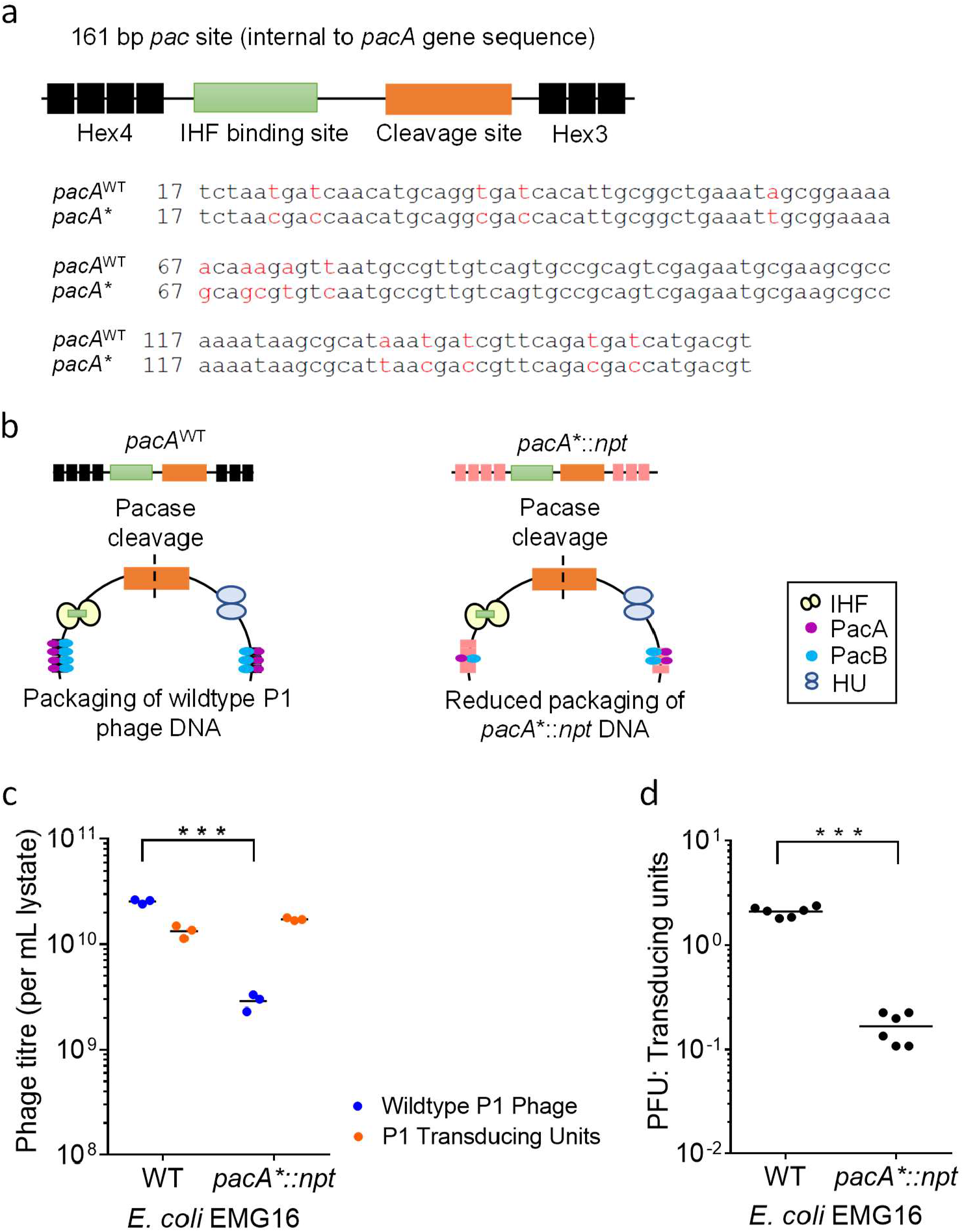
Genetic modification of *pac* site, within the *pacA* coding sequence of wildtype P1 bacteriophage genome, to reduce wildtype P1 phage DNA packaging. (**a**) Schematic diagram showing the minimal *pac* sequence of the P1 phage genome, situated within the *pacA* coding sequence. Methylation sites, consisting of hexameric repeats, HEX4 and HEX3 are shown in black, IHF binding site is shown in green and the *pac* cleavage site is shown in orange. Synonymous mutations were introduced onto the hexameric repeats to disrupt the binding of PacA and PacB. Comparison with wildtype *pac* DNA sequence is shown, with mutated DNA bases indicated in red. (**b**) Schematic diagram showing the processing of *pac* site, involving the binding of PacA and PacB to the hexameric repeats, as well as IHF and HU binding which was proposed to give a bent structure of the *pac* site. Pacase (both PacA and PacB) enzymatic activity then leads to cleavage of the *pac* site. Synonymous mutations introduced onto the hexameric repeats of the *pac* site lead to a reduction in pacase binding hence reducing the processing and packaging of the P1 mutant *pacA\**::*npt* P1 DNA. (**c**) Quantification of wildtype P1 phage (in blue) and P1 transducing units (in orange) titres of lysates prepared from wildtype (WT) and *pacA\**::*npt* EMG16 cells harbouring J72114 *cas9-*NT phagemid without spacer sequence(s) targeting *E. coli* or *S. flexneri* chromosomal sequences. (**d**) The ratio of wildtype P1 phage to P1 transducing units was calculated and plotted. Each data point represented a biological replicate and is the average of 4 technical repeats. Horizontal bars represent the group mean. The p-values (between wildtype and *pacA\**::*npt* lysates) were determined using a two-tailed unpaired t-test with significance defined by p < 0.05. p < 0.0005 between targeting phagemid and non-targeting phagemid treatments were shown as ***.

Overall, these results highlight an improvement in phagemid purity, with a significantly reduced ratio of wildtype P1 phage to phagemid for lysates prepared from *pacA\**::*npt* EMG16 mutant cell line (p < 0.0005, **Figure 3c**).

### Efficiency of *cas9* phagemid-induced cell lethality of *S. flexneri*

Results obtained so far support our hypothesis that the presence of a spacer sequence complementary to an *E. coli* chromosomal gene can lead to cell lethality via its Cas9 endonuclease activity. We next chose to test the delivery and properties of the J72114-*cas9* phagemid constructs in the context of *S. flexneri*, the causative agent of shigellosis (also called bacillary dysentery), with high rates of mortality and morbidity among children aged under 5 years in developing countries^21^. Spacer sequence mediated lethality effect of *cas9* phagemid on *S. flexneri* was first tested using *cas9* phagemids with spacer sequences designed to target 4 conserved virulence (and chromosomal) genes of *S. flexneri*: *sigA, pic, shiD, shiA* (refer to **Supplementary Table 3** for DNA sequences)^37,38,39,40^. To optimise the delivery of the *cas9* phagemid to a new host, a broad-host-range origin of replication pBBR1 was chosen for the J72114-*cas9* phagemid (**Figure 4a**). To validate the targeting efficiencies of the spacer sequences designed, we treated an avirulent strain of *S. flexneri* (strain 2a 2457O) using crude P1 *cas9* phagemid lysates prepared from wildtype *E. coli* P1 lysogen. These results showed that spacer sequence(s) targeting *shiA* or *shiD* genes gave the highest Cas9 mediated lethality effect on *S. flexneri*, with a ∼75-fold reduction in the recovered CFU as compared to that treated with non-(chromosomal) targeting *cas9-*NT phagemid (**Figure 4b**). Since the targeted virulence genes are not linked to survival of *S. flexneri* cells *in vitro*, the conditional lethality observed is likely to be due to DSB cleavage on the chromosome by the Cas9 endonuclease. These results indicate the functionality of the 4 spacer sequences in specific targeting of *S. flexneri* chromosomal genes for Cas9 mediated disruption and cell lethality.

**Figure 4:**
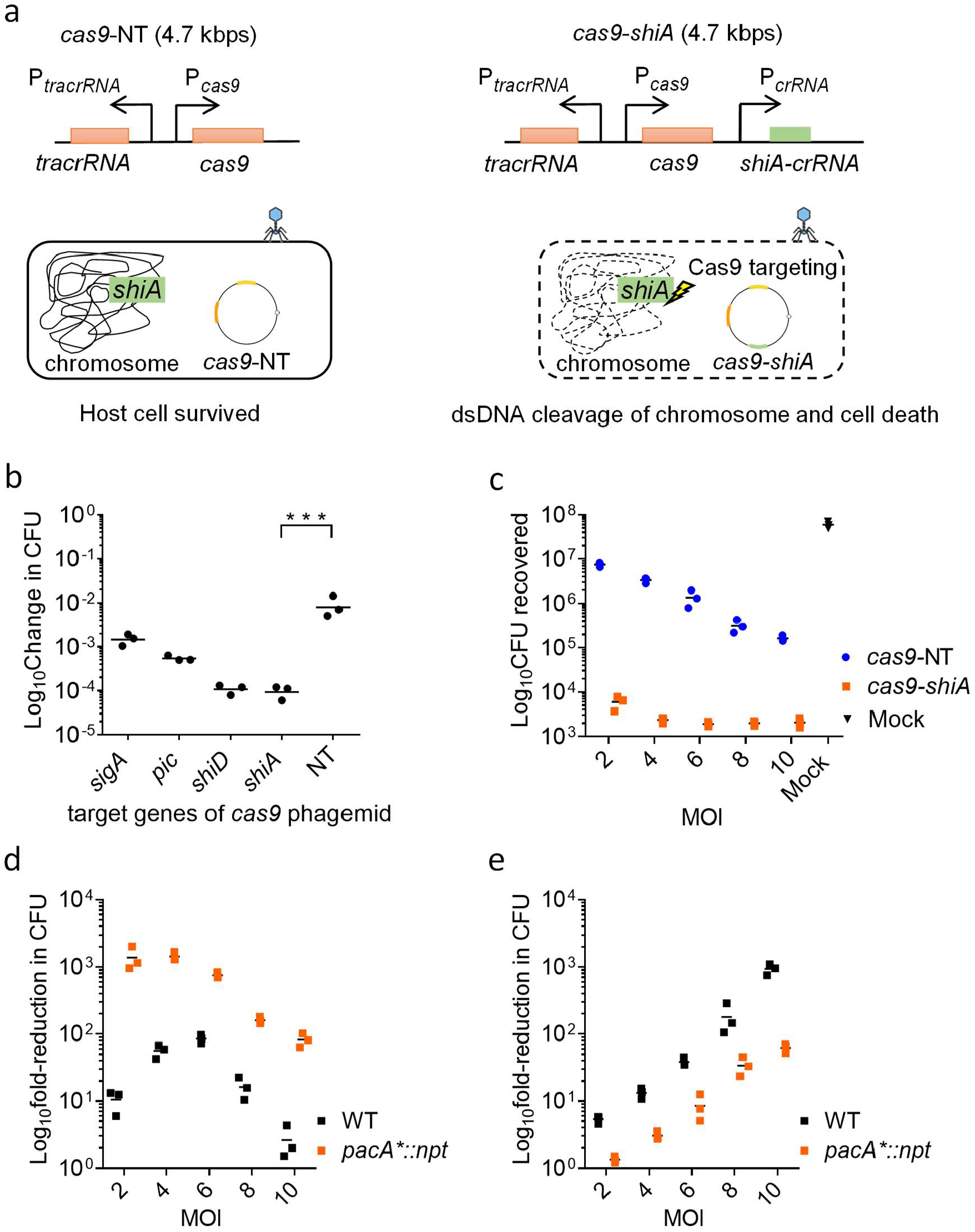
Cas9 mediated lethality of *S. flexneri* using chromosomal-targeting J72114 *cas9-shiA* phagemid. (**a**) Schematic diagram showing the *cas9* genetic construct of P1 J72114 phagemid, with spacer sequence targeting chromosomal gene(s) (i.e. *shiA*) of *S. flexneri* (in green). Upon transduction, the presence of *crRNA* with spacer sequence complementary to chromosomal genes of *S. flexneri* (ct-*crRNA*) would cause dsDNA cleavage of the chromosome via Cas9 endonuclease, leading to cell death. (**b**) J72114 *cas9* phagemid with spacer sequence(s) targeting *sigA, pic, shiD* and *shiA* chromosomal genes of *S. flexneri* 2a 2457O. Lysates were prepared from wildtype EMG16 cell line. Crude lysates were used for transduction of *S. flexneri* 2A 2457O, with an MOI of 10 wildtype P1 phage (equivalent to 5 transducing units) to 1 bacterial cell. Data were plotted as change(s) in CFU as compared to input CFU (approximately 1 × 10^7^ cells per reaction) used for infection. The p-value (between NT-phagemid and *shiA*-targeting phagemid treatment) was determined using a two-tailed unpaired t-test with significance defined by p < 0.05. p < 0.0005 between targeting phagemid and non-targeting phagemid treatments is shown as ***. (**c**) Spacer sequence mediated lethality of *S. flexneri* 5a MT905 cells with chromosomal-targeting *cas9-shiA* (in orange) and non-targeting, *cas9*-NT phagemids (in blue) lysates prepared from *pacA*^*\**^*::npt* EMG16 cell line. MOIs of 2.0, 4.0, 6.0, 8.0 and 10.0 were used. Data were plotted as changes in CFU as compared to input CFU (approximately 1 × 10^7^ cells per reaction) used for infection. The number of CFU recovered from mock infection was shown in black. (**d**) The Cas9 spacer sequence mediated lethality effect of *cas9*-*shiA* phagemid and (**e**) The non-spacer sequence mediated lethality effect of *cas9* phagemid lysate prepared from wildtype (WT, in black) and *pacA*\*::*npt* EMG (in orange) cell lines. MOIs of 2, 4, 6, 8, and 10 P1 transducing units to 1 bacterial cell for *pacA*\*::*npt* EMG lysates were used. For wildtype EMG16 lysates, MOIs of 2.0, 4.0, 6.0 and 8.0 and 10.0 wildtype P1 phage to 1 bacterial cell were used, and the wildtype P1 phage to P1 transducing units is approximately 2, for lysates prepared from wildtype EMG16 cell line. The *cas9-shiA* mediated lethality effect was quantified by measuring the reduction in CFU recovered after treatment with *cas9*-*shiA* and *cas9*-*NT* phagemids. The reduction in CFU recovered between *cas9*-NT phagemid treatment and the input cells used for infection (approximately 1 × 10^7^ cells per reaction) would show the non-spacer sequence mediated lethality effect of P1 phage lysates at all the 4 MOIs tested. Each data point represents a biological repeat and is the average of 4 technical repeats. Horizontal bars represent the group mean.

We next sought to validate the chromosomal-targeting efficiency of our *cas9-shiA* phagemid on a pathogenic strain of *S. flexneri*, strain 5a M90T, which is widely used as a paradigm for cellular microbiology studies and *in vivo* invasion assays^39^. The *cas9*-*shiA* phagemid was chosen for further transduction assays because it gave the highest spacer sequence-mediated lethality on *S. flexneri* 2a 2457O. P1 lysates of *cas9*-*shiA* phagemid, as well as its respective non-(chromosomal) targeting constructs (*cas9-*NT), were prepared from *pacA*\*::*npt* EMG16 cells, treated with PEG-6000 and used for transduction assays on *S. flexneri* 5a M90T. To identify the optimal dosage, a range of MOIs (2.0, 4.0, 6.0, 8.0 and 10.0) were used for the transduction assay. These results showed that an MOI value >2.0 gave the highest spacer sequence-mediated killing effect of *S. flexneri* cells, causing 3.8 × 10^3^ to 5.1 × 10^3^ reduction in *S. flexneri* CFU (p < 0.0005) after treatment with *cas9*-*shiA* phagemid (**Figure 4c**). The non-Cas9-mediated killing effect at MOIs of 2.0 to 4.0 reduced *S. flexneri* CFU by approximately 3.1-fold (p > 0.05) and 6.7-fold (p < 0.05) respectively, when compared to an input of 10^7^ *S. flexneri* CFU. We determined that an MOI value of ∼4.0 is optimal, considering that a further increase in MOI would lead to an increase in the non-spacer sequence mediated lethality effect of lysates without significant changes to the Cas9 mediated lethality effect (**Figure 4c**). We did not recover any chloramphenicol-resistant *S. flexneri* CFU after treatment with *cas9-shiA* phagemid, thus validating the efficiency of its spacer sequence mediated cell lethality.

To compare the effect of *pacA*\*::*npt* mutation on improving the quality of lysate, results were compared to that of lysates prepared from wildtype EMG16 cells. Lysates prepared from wildtype EMG16 cells required an MOI of approximately 6.0 for a maximum Cas9 mediated lethality effect of ∼100-fold reduction in CFU (**Figure 4d**), which was accompanied by a non-spacer sequence mediated killing effect of approximately 40-fold reduction in CFU (**Figure 4e**) (p < 0.0005 for comparison of CFU recovered between *cas9*-NT and *cas9*-*shiA* treatment). The maximum Cas9-mediated lethality effect of *pacA*\*::*npt* lysates can be achieved at a lower MOI of approximately 4.0, thus giving an overall lower non-spacer sequence killing effect, as compared to that observed for lysates prepared from wildtype EMG16 cells (**Figure 4d, 4e**).

These results demonstrate the specificity and efficiency of our *cas9-shiA* phagemid in killing *S. flexneri* 5a M90T cells *in vitro*. We concluded that *pacA*\*::*npt* lysates could achieve its maximum Cas9 mediated lethality effect at a lower MOI as compared to lysates prepared from wildtype EMG16 cells, hence giving a lower non-spacer sequence killing effect of phagemid lysates on *S. flexneri* cells.

### Validating the efficiency of the P1 *cas9* phagemid system *in vivo* to control lethal *S. flexneri* infection in zebrafish larvae

We next sought to establish whether our P1 *cas9* phagemid system could clear *S. flexneri* infection *in vivo*. A variety of studies have shown that zebrafish larvae are susceptible to *S. flexneri* infection, with key aspects of the human disease being replicated in this model^41,42,43^. Zebrafish larvae are recognised as highly versatile for studying innovative treatments against *S. flexneri* infection^43,44^, for example, clearance of drug-resistant *S. flexneri* infection *in vivo* has been achieved via the injection of predatory bacteria *Bdellovibrio bacteriovorus*^42^. To assess the spacer sequence-mediated killing effect of our P1 *cas9* phagemid, P1 phage lysates of *cas9*-*shiA* and *cas9-*NT phagemid (without chromosomal-targeting spacer sequence) were injected into the hindbrain ventricle of zebrafish larvae at 2 days post-fertilisation, following the administration of a lethal dose (∼8000 CFU) of *S. flexneri* 5a M90T. We observed a ∼10-fold reduction in *S. flexneri* CFU after treatment with *cas9*-*shiA* phagemid at 6 h post-infection, compared to that of non-targeting *cas9-*NT phagemid treatment (p < 0.005, **Figure 5a**). This was accompanied by a ∼20% in survival rate of zebrafish larvae (p < 0.005) (**Figure 5b**). Injection with either the *cas9*-*shiA* or c*as9*-NT phagemid alone without *S. flexneri* infection did not lead to morphological defects or reduced viability of zebrafish larvae (**Supplementary Figure 3**). Overall, these results demonstrate the efficiency of *cas9*-*shiA* chromosomal-targeting phagemid in reducing *S. flexneri* bacterial load *in vivo* and improving the survival of the infected host (**Figure 5c**). Considering the non-replicative nature of the P1 phagemid, our results highlight the potential use of *cas9* phagemids as a safe and efficient therapeutic agent *in vivo*.

**Figure 5:**
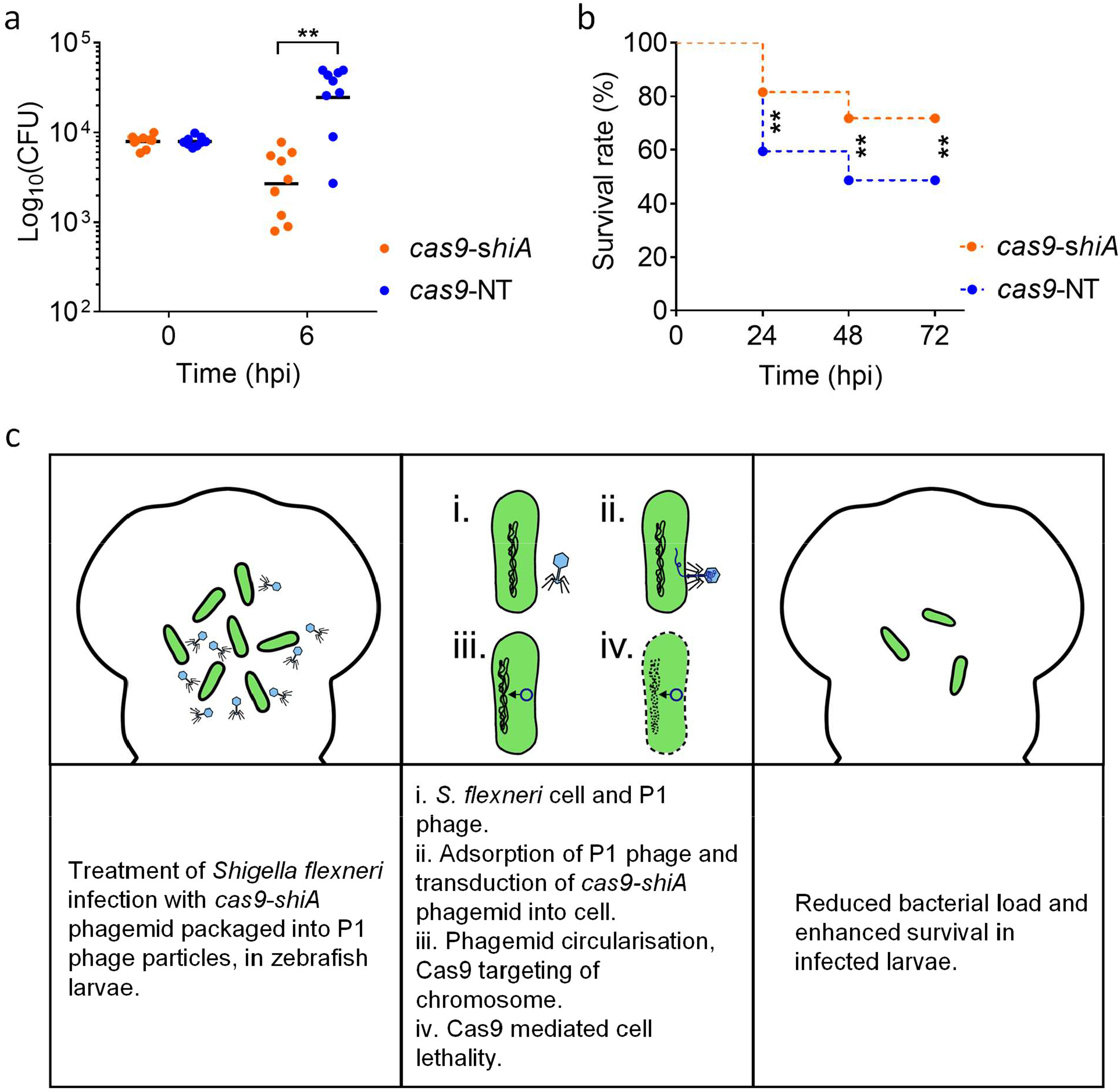
Spacer sequence mediated lethality of *Shigella flexneri* 5a MT905 *in vivo* using chromosomal-targeting *cas9*-*shiA* phagemid. (**a**) Enumeration of *S. flexneri* 5a M90T CFU at 6 h post-infection. n = 9 (targeting); 9 (empty) larvae (cumulated from 3 independent experiments). (**b**) Measurement of survival rate of infected zebrafish larvae at 24, 48 and 72 h post-infection (hpi), treated with *cas9*-*shiA* phagemid (blue) or with the non-targeting *cas9*-NT phagemid (red). n = 71 (targeting); 74 (empty) larvae (cumulated from 3 independent experiments). Differences in bacterial load were tested using an unpaired t-test on Log10-transformed data while differences in survival were tested using a Log-rank (Mantel-Cox) test. p < 0.05 is considered statistically significant. p < 0.005 between targeting phagemid and non-targeting phagemid treatments were shown as **, at the time points indicated on the graph. (**c**) Schematic diagram showing the reduction of *S. flexneri* infection bacterial load via the spacer sequence mediated lethality effect of *cas9*-*shiA* phagemid. The reduced bacterial load would promote survival of infected larvae.

## DISCUSSION

In this study we demonstrate P1 phagemid-based delivery of chromosomal-targeting *cas9* genetic construct into *E. coli* and *S. flexneri* cells. We establish protocols that give a high phagemid titre, and introduce the use of *pacA*\*::*npt* EMG16 cell line, which gives improved phagemid purity of lysates. We show efficient killing of *S. flexneri* cells with *cas9* phagemid in the presence of spacer sequences complementary to chromosomal gene(s) of *S. flexneri*. Finally, treatment of *S. flexneri* infected zebrafish larvae with chromosomal-targeting *cas9-shiA* phagemid significantly reduced bacterial burden and improved host survival.

We chose the P1 bacteriophage to deliver our phagemids into *E. coli* and *S. flexneri* due to its 1) ability to package large-sized DNA (∼100 kbps), 2) high transduction efficiency among Gram-negative *Enterobacteriaceae*, 3) ability to lysogenise after transduction, and 4) *in trans* induction of single gene expression, *coi*, which can promote lytic stage replication^25^. Using the P1-phagemid system with arabinose-inducible *coi* gene expression, our results are consistent with observations from previous studies, showing a transduction efficiency of at least 7 × 10^8^ transducing units per mL lysate used on *E. coli*^24,25^. We observed a significantly lower transduction efficiency of the P1 phagemid for *E. coli* TOP10 cell, and such reduced transduction efficiency was previously reported on several *recA-* strains of *E. coli*^24^. Previous studies suggested that only a subset of P1 phage particles contain Cre recombinase while the rest of the P1 phage DNA relies on the host cells’ homologous recombination system, such as RecA and RecBCD mediated homologous recombination of Chi sites, for circularisation of linear genomic DNA^45,46^. Chi sites are over-represented in the genome of the P1 bacteriophage, and the *cin* gene sequence contains 2 of the 50 identified Chi sites which may have improved the transduction efficiency of our P1 phagemid^46^.

Our results demonstrate that the P1 phagemid is efficient in delivering the *cas9* genetic constructs into both *E. coli* and *S. flexneri*. We show that the presence of spacer sequences complementary to the targeted chromosomal gene(s) of *E. coli* and *S. flexneri* yielded 2 to 3 log reduction in bacterial CFU *in vitro*, which is comparable with the Cas9-chromosomal-targeting effect reported in previous studies^5,6,17,33^. The efficiency of the *cas9*-*shiA* targeting phagemid in reducing *S. flexneri* bacterial load *in vivo* was demonstrated as early as 6 h post-infection, with a ∼10-fold reduction in CFU, thus leading to a ∼20% increase in the survival rate of infected larvae. The reduced spacer sequence mediated lethality effect observed *in vivo* as compared to that observed *in vitro* may be due to differences in experimental conditions (i.e. temperature, duration of infection, environmental conditions) and the lifecycle of *S. flexneri* infection *in vivo*. We hypothesise that the P1 bacteriophage unable to cross the host cell membrane, and thus the primary target cells of the phagemid are the extracellular pool of *S. flexneri*. Previous studies showed that repeated administration of phages enhanced clearance of bacterial infections *in vivo*, which might improve the performance of our *cas9* phagemid^47,48,49^. However, we did not distinguish whether the surviving *S. flexneri* colonies recovered after *shiA*-targeting phagemid lysates treatment were P1-resistant colonies and/or escape mutants of the Cas9 chromosomal-targeting effect. If the surviving *S. flexneri* are resistant to P1 infection, repeated administration of phagemid lysates may not improve the overall Cas9-mediated killing of bacteria. The factor(s) which may be associated with resistance against phage infection and/or the Cas9 chromosomal-targeting system could be identified by genomic sequencing of surviving *S. flexneri* colonies. These data could potentially provide insights into ways of improving our current P1 phagemid system, such as engineering of P1 tail fibre to overcome phage resistance, the use of multiple chromosomal-targeting guide RNAs and/or the use of other RNA-guided endonucleases to enhance the antimicrobial effect. In addition, our study lacks pharmacokinetic analysis to investigate the interactions between P1 phage and the host immune response, which could dictate the efficiency of our P1 phage-based delivery method. Future assessment of the interactions between P1 phage and the host immune system may provide solutions to enhance the stability of P1 phage, which could improve clearance of infection *in vivo*.

Given the non-replicative nature of P1 *cas9* phagemid, a higher MOI is required for a significant spacer sequence mediated lethality effect. However, our results showed that an increase in MOI would also lead to an increase in the general cytotoxicity effect of phagemid lysates on *E. coli* and *S. flexneri*. We demonstrated that this could be mitigated via PEG-6000 treatment of lysates and the genetic modification of the *pac* site on P1 genome, by reducing the recovery of wildtype P1 phage and its DNA packaging, respectively. The improved phagemid purity allowed higher spacer-specific lethality at a lower MOI. Given the “headful” DNA packaging mechanism of P1 that requires ∼100 kbp of DNA subtrate for phage maturation, as well as the ability of P1 in packaging DNA substrate without *pac* sequence, production of pure P1 phagemid transducing particles may be challenging^50,51^. Although the well-studied M13-based phagemid system yields a higher titre of pure transducing particles compared to the P1 phagemid system, M13 adsorption requires the tips of F-pili, which restricts its host range to F^+^ cells only^52,53^. This might limit the efficiency of phagemid delivery, especially in targeting clinical isolates of *S. flexneri*, when compared to P1 phage transduction which has a broader host range. Tridgett et al., (2021) demonstrated the production of pure cosmid transducing particles, using a mutant P2 lysogen that has its DNA packaging site, *cos*, replaced with P4 *δ* and *ε* gene sequences^54^. However, our preliminary results suggest a lower transduction efficiency of cosmid DNA into *S. flexneri* 5a M90T by P2 when compared to P1 infection, despite both phages having broad host ranges (data not shown). We are currently developing a P4 cosmid system which could produce pure cosmid transducing particles, using the mutant strain of P2 lysogen described by Tridgett et al.^54^ Replacing the host range determining region of P2 tail fibre with that of P1 may improve the transduction efficiency of cosmid DNA into *S. flexneri* 5a M90T. Comparisons between P1 phagemid lysates and P4 cosmid lysates should provide insights into the effects of using pure cosmid transducing particles on the non-Cas9 killing of *S. flexneri*.

The variation across *S. flexneri* serotypes, which change regionally, is likely to complicate the development of an effective and broadly-protective vaccine against *S. flexneri* infection^55,56^. The versatility of the CRISPR Cas9 system allows seamless reprogramming of the endonuclease to target conserved chromosomal DNA sequences and/or virulence factors encoded by *S. flexneri*, by modifying its CRISPR guide RNA spacer sequence. Although our results indicate Cas9 killing of *S. flexneri* with 4 different chromosomal-targeting guide RNAs, a wider panel of spacer sequences may be useful to identify target sites improving the Cas9-mediated killing of *S. flexneri*. We demonstrate that P1 phage can transduce phagemid DNA into both *S. flexneri* serotypes 2a and 5a, highlighting the great potential of P1 as a universal *S. flexneri* targeting strategy, as the targets can be selected to be conserved across *S. flexneri* serotypes. We have observed that the transduction of phagemid DNA by P1 with its alternative S’ tail fibre is not significantly affected by mutations in the O-antigen modification genes of *S. flexneri* 2a 2457O and 5a M90T (data not shown). *S. flexneri* serotypes, except serotype 6 (Sf6), share the same O-antigen backbone^57^, therefore P1(S’) could potentially be exploited to transduce phagemid DNA into other serotypes of *S. flexneri*. We also assessed P1(S’) transduction on serotype 2b, which yielded a higher number of phagemid transductants when compared to P1(S) infection (data not shown). In the future, it will be interesting to assess P1 transduction efficiency on the remaining *S. flexneri* serotypes, to determine if the P1 *cas9* phagemid can provide broad and efficient targeting of the bacteria.

In summary, combining CRISPR-Cas9 sequence-specific bacterial targeting with P1 bacteriophage-based delivery has great potential to be used as a supplement to conventional antibiotics for the treatment of antibiotic-resistant bacterial infections. As demonstrated in this study, the genetic modification of P1 bacteriophage and the incorporation of the CRISPR-Cas9 system in the form of phagemid is useful for targeting clinically relevant Gram-negative *Enterobacteriaceae*.

## MATERIALS AND METHODS

### Bacterial and phage strains, plasmids/phagemids and growth media

Strains of *E. coli* and *S. flexneri* used for this study are listed in **Supplementary Table 1**, with a full description of strain modification (if any), as well as the purpose of each strain used in this study. Plasmids/phagemids used in this study are listed in **Supplementary Table 2** and were constructed using Gibson assembly. DNA sequences of constructs are listed in **Supplementary Table 3**. Bacterial cells were cultured in Luria–Bertani medium (LB) or phage lysis medium (PLM; LB containing 100 mM MgCl2 and 5 mM CaCl2), while SM buffer (50 mM Tris-HCl, 8 mM MgSO4, 100 mM NaCl, pH 7.5), was used for P1 bacteriophage manipulation as stated in previous studies of P1 bacteriophage^24,25^. Concentrations of antibiotics used were 50 μg mL^−1^ for ampicillin and kanamycin, and 25 μg mL^−1^ for chloramphenicol. All chemical reagents used were analytical grade and purchased from Sigma-Aldrich.

### Bsa1-cloning of chromosomal-targeting protospacer sequences

The CRISPR Cas9 construct used in this study was derived from a pCas9 plasmid (Addgene plasmid #42876), containing the *cas9* gene under a constitutive promoter, a *trans*-activating CRISPR RNA (*tracrRNA*) and a CRISPR guide RNA (*crRNA*). The *crRNA* sequence contains two BsaI sites that allow molecular cloning of a protospacer sequence. 20 bps protospacer sequences targeting the *npt* and *S. flexneri* chromosomal sequence(s) with a NGG protospacer adjacent motif (PAM) were designed using the CHOP-CHOP web tool^26^. Primer pairs having a protospacer sequence were designed to have compatible ends for its annealing into the Bsa1-digested sites of the *crRNA* sequence. The design, annealing and cloning of primer pairs into the *crRNA* sequence were performed as described by Jiang et al.,(2013)^27^. Restriction digestion of phagemid DNA was carried out using BsaI-HF®v2 (NEB), following the manufacturer’s protocol. Primer pairs used for protospacer sequences cloning are listed in **Supplementary Table 4**.

### Phage lysates preparation

Phage lysates were prepared using a previously established protocol, with some modifications^25^. Briefly, *E. coli* EMG16 harbouring the P1kc lysogen (P1 is used throughout the text instead), were chemically transformed with the J72114-*cas9* phagemids, using a standard protocol for heat-shocked transformation of *E. coli* cells. An overnight culture of transformed *E. coli* P1 lysogen was diluted 1/100 in fresh PLM media, and cultured for 1 h at 37°C. Cell lysis was induced via the addition of L-arabinose (final concentration of 13 mM). Cell lysis was defined as the presence of debris, and the clearance of bacterial culture, which happened at approximately2 h post-induction with 13 mM L-arabinose. Chloroform (final concentration of 2.5%) was added to lysates and cultures were shaken for 30 min, to aid in thorough cell lysis and lysate sterilisation. Cultures were vortexed, lysates were clarified via centrifugation at maximum speed (16,000 g for 3 min), and the supernatant was collected. The supernatant was then passed through a 0.22 μm syringe filter (Millipore) to remove the remaining cell debris. At this stage, the lysates prepared were identified as “crude lysate”, and stored at 4°C.

Our preliminary results suggested that lysates produced from *E. coli* strain NCM3722 harbouring P1 contained a higher number of transducing units as compared to that of strain EMG16 (**Supplementary Figure 4**). For the preparation of phage lysates using NCM3722 P1, overnight culture of *E. coli* NCM3722 cells was diluted in fresh LB medium and cultured at 37°C until an OD600 of approximately 1.0. Cells were 10-fold concentrated in PLM medium. Crude lysates prepared from *E. coli* EMG16 cells were used to transduce NCM3722 cells (refer to transduction section of Materials and Methods). Cells were plated onto LB agar with kanamycin and chloramphenicol which selects for both the *pacA*\*::*npt* P1 and the J72114-phagemids respectively, and plates were incubated at 37°C for at least 16 h. For lysates prepared from wildtype P1 lysogen, transduced NCM3722 cells were plated onto LB agar with chloramphenicol only. Colonies were picked, and PCR reaction(s) were carried out using primers that anneal specifically to genes of the P1 genome, such as *lpa*. The same protocol for arabinose induction of cell lysis mentioned above was used for making lysates from NCM3722 cells harbouring both P1 and the J72114-phagemids.

### PEG-6000 treatment of phage lysate

Treatment of crude lysates with PEG-6000 was carried out based on protocols established in previous studies but with slight modification^28,29^. Briefly, crude lysates prepared were first treated with NaCl to a final concentration of 0.33 M, and incubated on ice for 1 h. This step was to ensure the precipitation of debris and proteins in the lysates. Lysates were centrifuged at 5,000 g for 50 min, the supernatant was collected and further treated with PEG-6000 to a final concentration of 4%, at 4°C overnight. The use of 4% PEG-6000 was justified by our results which showed a significant reduction in the amount of wildtype P1 phage, and a substantial number of transducing units recovered from the lysates (data not shown). After overnight incubation with PEG-6000, phage precipitate was spun down at 5,000 g for 1 h at 4°C. The supernatant was removed and the phage precipitate was resuspended in an appropriate volume of SM buffer, i.e. 50X concentration of the phage particle would require resuspending the phage precipitate with 1 mL of SM buffer for 50 mL of lysate. Lysates were filtered through a 0.22 μm syringe filter (Millipore), and further washed or concentrated using Amicon Ultra MWCO 100 kDa centrifugal filter units (Millipore). Lysates were then stored at 4°C.

### Quantification of plaque-forming units (PFU)

Plaque assay was carried out to estimate the population of wildtype P1 phage in lysates, using previously established protocols^30^. Briefly, stationary phase culture of naïve *E. coli* host strain, NCM3722, was diluted in fresh PLM broth by 1/100, cultured at 37°C with shaking until an OD600 of 0.5, which took approximately 2.5 h. From our preliminary results, phage lysates prepared from wildtype and *pacA*\*::*npt* P1 *E. coli* lysogen would have to be diluted to 10^7^ and 10^6^, respectively, to give a reasonable amount (50 to 200) of PFU. 1 mL of cell suspension was added to 100 μL of diluted phage lysate, vortexed, and incubated at 37°C with shaking for 10 min. Cells and phage mixture were added to 3 mL of melted top agar (LB medium with 0.6 % agar), then poured immediately onto bottom agar (LB medium with 1.5 % agar). Agar plates were dried at room temperature for 10 min, then incubated at 37°C for at least 16 h, before the enumeration of PFU.

### Quantification of transducing units/phagemid

Transduction assay was carried out based on protocol established by a previous study with slight modifications^25^. Instead of using a stationary phase culture for transduction, an overnight culture of the indicator *E. coli* strain NCM3722, was first diluted in fresh LB broth by 1/100 and cultured at 37°C with shaking until an OD600 of 0.5. Cells were spun down at 3,000 g for 5 min, concentrated 10-fold in fresh PLM buffer, providing approximately 10^8^ cells per 100 μL of bacterial suspension for transduction. An equal volume of diluted (10^1^ to 10^2^ dilution factor) phage lysate was mixed with the resuspended cells, and phage adsorption was allowed for a maximum of 30 min at 37°C with shaking. SOC with 10 mM sodium citrate was added to the cell and phage lysate mixture for recovery of cells, expression of antibiotic resistance marker, and quenching of further phage infection via citrate interaction with free calcium ions needed for phage adsorption. SOC recovery was carried out at 37°C for 1 h with shaking. Serial dilutions of recovered cells were performed, and spotted onto plain LB agar, as well as LB agar supplemented with 25 μg/mL of chloramphenicol. Agar plates were incubated at 37°C for at least 16 h, before the enumeration of chloramphenicol-resistant colonies. Transducing efficiency would be defined by the percentage of chloramphenicol resistant colonies against total CFU recovered on plain LB agar.

### *pacA* genetic modification

Lambda red recombineering technique was used for genetic manipulation of *pacA* gene of P1 genome. The template for recombination was designed to have homology arms complementary to the 3’-end of *lpa* gene and the 5’-end of *pac* sequence (**Supplementary Tables 3**). Synonymous mutations were introduced via codon optimisation of the *pac* site to disrupt the hexameric repeats. A kanamycin resistance cassette was included in the *pacA* modification template for lambda red recombination as a selection marker for positive clones, which would be integrated into the intergenic region between *lpa* gene and LpPac promoter. The *pacA* modification template was first cloned into an empty plasmid with pSC101 origin of replication, via Gibson assembly and sequenced verified. Linear dsDNA substrate used for lambda red recombination was produced via PCR, followed by gel extraction of PCR product. Lambda red recombineering was carried out based on previously established protocol^31^. Briefly, wildtype *E. coli* EMG16 P1 lysogen was transformed with pKD46 plasmid. Stationary phase culture of the transformed cells was grown in fresh LB at a dilution factor of 100, at 30°C with shaking, until an OD600 of 0.35. 0.65 M L-arabinose was added to the culture to induce the expression of lambda Red genes (*exo, bet, gam*), and cultured for a maximum of 30 min at 30°C, with shaking. Cells were chilled on ice for 40 min and made electrocompetent, using standard protocols for preparing electrocompetent cells. 100 ng of dsDNA substrate was electroporated into the cells, followed by growth in SOC at 30°C with shaking for 2 h. Cells were plated onto LB agar supplemented with 50 μg/mL of kanamycin, and incubated at 37°C for at least 16 h. Colony PCR was carried out on colonies recovered, using primer pairs that a) anneal to the junction of integration to verify the correct insertion of the modification template (refer to **Supplementary Figure 5**) and b) are complementary to the modified nucleotide bases to select for colonies that retain the mutations to the *pac* site. PCR products (using primers annealing to the junction of integration) of the correct size were excised and sequence-verified. Positive clones were re-streaked onto new LB agar supplemented with 50 μg/mL of kanamycin, incubated at 37°C for at least 16 h, and this re-streaking process was repeated for at least 3 generations to ensure homogeneity in the bacterial colony. The kanamycin resistance cassette was retained in the mutant cell line which provided a selectable marker for the *pacA*\*::*npt* P1 lysogen.

### *E. coli* MC1061::*npt* and *S. flexneri* chromosomal-targeting assay

Stationary phase culture of naïve host cells *E. coli* MC1061::*npt* were sub-cultured in fresh PLM at a dilution factor of 100 at 37°C until an OD600 of 0.35 is reached. Cells were then diluted to reach an OD600 of 0.1 in fresh PLM, which gave approximately 1 × 10^8^ cells per mL culture. 100 μL of the diluted culture, giving 1 × 10^7^ cells, was mixed with an equal volume of phage lysate diluted to the intended MOI. Phage adsorption was allowed for 30 min with shaking at 37°C. Cells were then recovered and further phage infection was quenched by the addition of SOC with 10 mM sodium citrate, for 1 h at 37°C with shaking. Serial dilutions of cells were made and spotted onto plain LB agar and/or LB agar with 25 μg/mL of chloramphenicol or 50 μg/mL of kanamycin. To enumerate the input used for chromosomal-targeting assay, 100 μL of the diluted culture was combined with an equal volume of SM buffer, followed by the addition of SOC with 10 mM sodium citrate then plated onto plain LB agar and/or LB agar with 50 μg/mL of kanamycin. Mock infected cells were treated the same as input cells, but with an additional 1.5 h shaking at 37°C, similar to cells treated with phage lysates. Agar plates were incubated at 37°C for at least 16 h before enumeration of CFU. The number of recovered CFU was then normalised to that of input cells, except for those plated on LB agar with chloramphenicol, whereby data was normalised to the number of CFU recovered after treatment with phagemids without *npt*-targeting spacer sequence, since input cells would not have the phagemid hence not chloramphenicol resistant.

### Zebrafish larvae model for *in vivo S. flexneri* infection

Animal experiments were performed according to the Animals (Scientific Procedures) Act 1986 and approved by the Home Office (Project licenses: PPL P84A89400 and P4E664E3C). Protocols are in compliance with standard procedures as reported at zfin.org. Unless specified otherwise, eggs, embryos and larvae were reared at 28.5°C in 0.5 X E2 medium supplemented with 0.3 μg/ml methylene blue. Injections were performed under anaesthesia, obtained by supplementing the medium with buffered 200 μg/ml tricaine.

GFP fluorescent and carbenicillin resistant *Shigella flexneri* 5a M90T was prepared for injections as in Torraca et al. 2019^32^. Bacteria were suspended to ∼8000 CFU/nl in PBS containing 2% polyvinylpyrrolidone and 0.5% phenol red. 1 nl of the bacterial suspension was microinjected in the hindbrain ventricle of 2 days post-fertilisation (dpf) zebrafish larvae. At 45 min post-infection, infected larvae were injected with a phagemid suspension (3 nl of a phagemid solution in SM Buffer + 5 mM CaCl2, corresponding to ∼10^4^ P1 transducing units).

Following phagemid delivery, larvae were incubated at 32.5°C. Survival rate was recorded at 24, 48 and 72 h post-infection. Bacterial enumeration from zebrafish was performed at 0 and 6 hpi by mechanical disruption of infected larvae in 0.4% Triton X-100 and plating of serial dilutions onto Congo red-tryptic soy agar plates containing 100 ug/ml carbenicillin.

### Statistical analysis

All experiments were carried out with at least 3 biological repeats and 4 technical repeats. Calculations of results were performex in Excel (Microsoft, Redmond, WA, USA), or Graphpad Prism 6. Graphpad Prism 6 was used to generate graphs. Student’s t-test (unequal variance, 2-tailed) or one-tailed ANOVA (for **Figure 1c**) were carried out to determine statistical significance. Data are expressed as means. For zebrafish experiments, differences in bacterial load were tested using an unpaired t-test on Log10-transformed data while differences in survival were tested using a Log-rank (Mantel-Cox) test. p < 0.05 is considered statistically significant. Stars on graphs represent p-values for statistically significant comparisons, with * denotes p < 0.05, ** denotes p < 0.005, *** denotes p < 0.0005 and ns representing p > 0.05 or non-significance.

## Data availability

All data in the main text and the supplementary materials are available from the corresponding author upon reasonable request.

## ACKNOWLEDGEMENTS

This work was supported by the Bill and Melinda Gates Foundation under the Grand Challenges Explorations grant (OPP1139488), and UK Research and Innovation Future Leaders Fellowship [MR/S018875/1]. V.T. is supported by an LSHTM/Wellcome Trust Institutional Strategic Support Fund (ISSF) Fellowship (204928/Z/16/Z). S.L.M. is supported by a Biotechnology and Biological Sciences Research Council LIDo PhD studentship (BB/T008709/1). Work in the S.M. laboratory is supported by a European Research Council Consolidator Grant (grant agreement No. 772853-ENTRAPMENT), Wellcome Trust Senior Research Fellowship (206444/Z/17/Z), and the Lister Institute of Preventive Medicine.

## AUTHOR CONTRIBUTIONS

B.W. conceived and supervised the study. B.W., Y.H., V.T., R.B. and S.M. designed the experiments. Y.H. and R.B. performed the experiments related to the P1 phagemid design, construction, production and characterisation in bacterial culture and P1 phage genome engineering. V.T. and S.L.M. performed zebrafish infection experiments. Y.H., V.T., R.B., J.F. and D.O. performed data analysis. All authors took part in the interpretation of results and preparation of materials for the manuscript. Y.H., S.M. and B.W. wrote the manuscript with input from all authors.

