## Supplementary Information for "P1 bacteriophage-enabled delivery of CRISPR-Cas9 antimicrobial activity against *Shigella flexneri*"

**Table of Contents**

Supplementary Figures 1-5

Supplementary Tables 1-4

Supplementary References

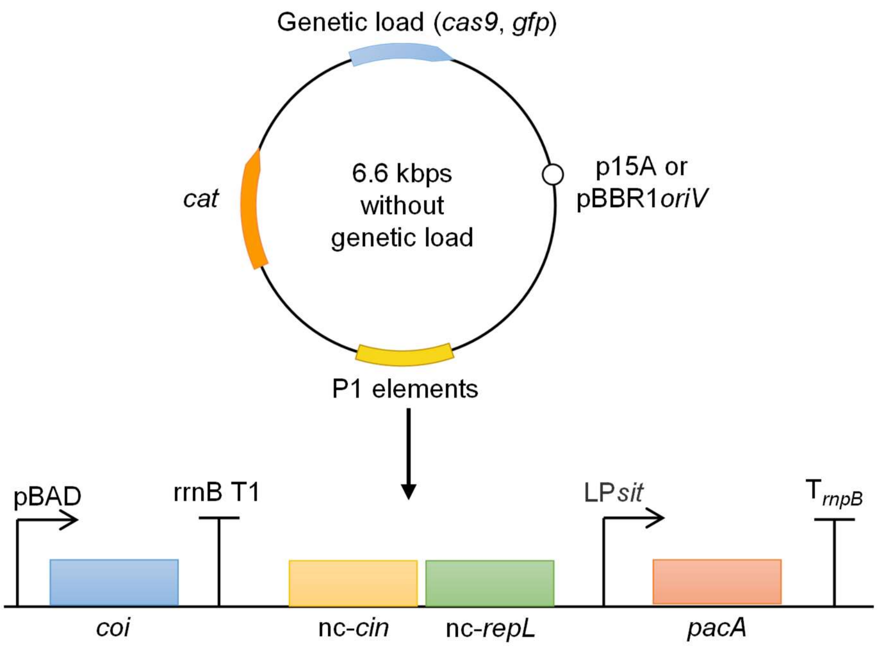


**Supplementary Figure 1: Schematic diagram showing the design of the J72114 phagemid.** The P1-based elements of our J72114 phagemid series are *coi*, *pacA*, non-coding copy of *repL* and *cin* gene sequences[1,2]. *coi*, whose gene product acts as a repressor antagonist of c1 repressor, hence acting as the switch that induces the lytic stage replication of P1 bacteriophage[3]. The *coi* gene expression of the BBa_J72114 phagemid was placed under the regulation of arabinose-inducible promoter, $P_{BAD}$, thus allowing *trans*-activation of the phagemid packaging into phage particles in the presence of arabinose. A non-coding copy of the P1 *repL* gene, which contains the P1 lytic stage origin of replication, $ori_L$, was included in the BBa_J72114 phagemid, allowing the lytic replicase generating phagemid DNA in the linear concatenated form required for packaging into the P1 virion during virus assembly[1,2]. A copy of the P1 *pacA* gene is also included under the control of the late lytic promoter $LP_{sit}$. The hexamer repeat motif of *pacA* is recognised by the P1 pacase, which cleaves at this site and brings the concatemeric DNA into an empty P1 capsid for packaging[1,2]. Finally, a non-coding copy of *cin*, a recombinase involved in expanding the P1 host range through tail-fibre variation, is included to increase phagemid packaging efficiency via an unknown mechanism[2]. The p15a origin of replication was swapped to the broader host spectrum pBBR1 for the phagemids used for chromosomal targeting of *S. flexneri.* All of our J72114 phagemid series contain a copy of *cat* gene, which confers transduced bacterial cells with chloramphenicol resistance.

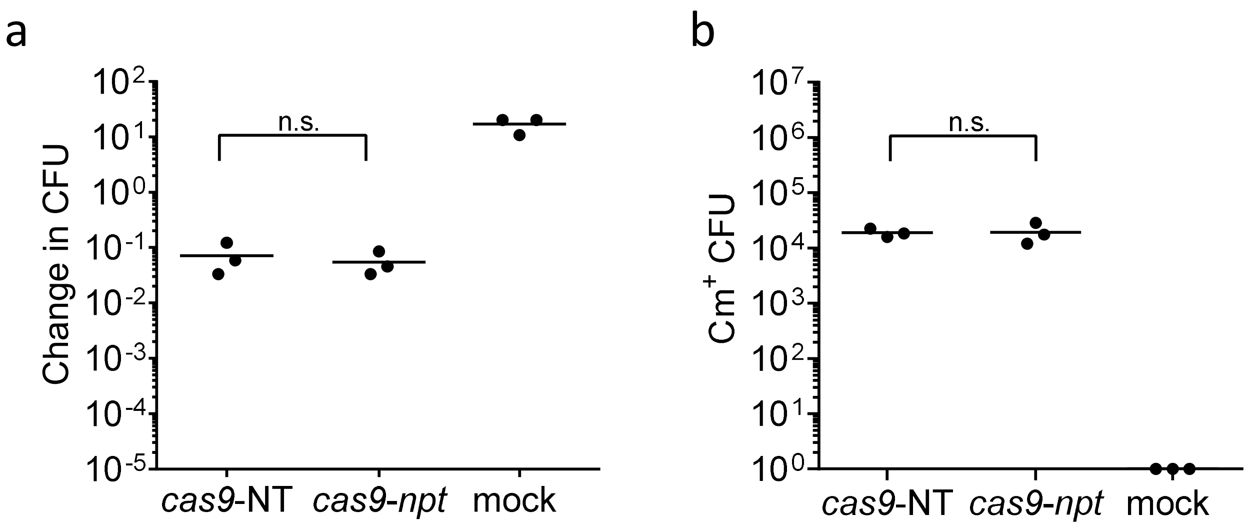


**Supplementary Figures 2: No significant Cas9-mediated lethality effect in *E. coli* K12 MC1061 cells without the chromosomal *npt* gene.** (**a**) Serial dilutions of transduced *E. coli* K12 MC1061 cells were plated onto plain LB agar. Data were plotted as change(s) in CFU as compared to input CFU (approximately $10^7$ cells per reaction) used for infection. (**b**) Quantification of chloramphenicol-resistant CFUs recovered, after treatment with *cas9*-NT or *cas9-npt* phagemid lysates. Each data point represents a biological replicate and is the average of 4 technical repeats. MOI of 10 wildtype P1 phage (equivalent to 5 transducing units) to 1 bacterial cell was used for all infections. Data was represented in the form of mean. The p-values (between non-targeting and targeting phagemid treatments) were determined using a two-tailed unpaired t-test with significance defined by $p < 0.05$. n.s. represents no significant difference(s).

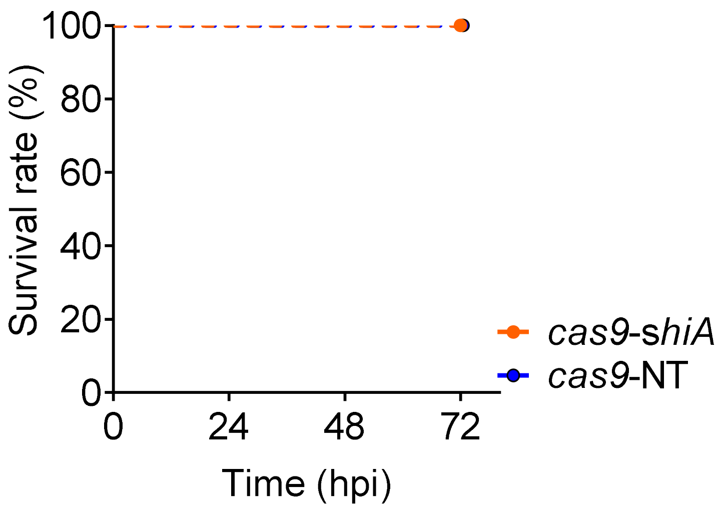


**Supplementary Figure 3**: **Injection of *cas9* phagemids does not affect zebrafish viability.** Survival curve of zebrafish larvae injected with ~$10^4$ P1 transducing units of *cas9-shiA* phagemid (blue) or *cas9*-NT phagemid (orange) without administering a lethal dosage of *S. flexneri*, incubated at 32.5°C for 72 hours post-infection (hpi). Without *S. flexneri* infection, zebrafish larvae showed a 100 % survival rate at all hpi after *shiA*-targeting and non-targeting phagemid treatments, suggesting that both the Cas9-killing effect and lysates cytotoxicity were specific towards *S. flexneri*.

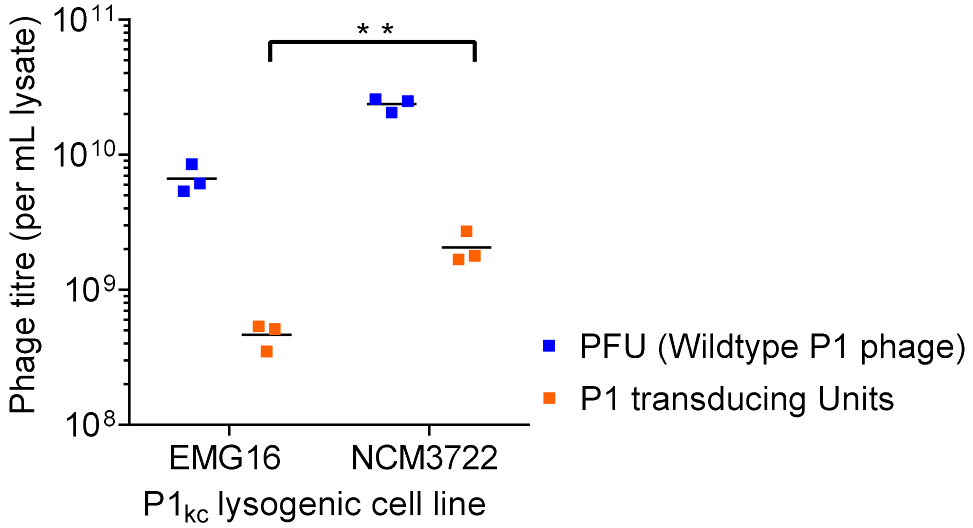


**Supplementary Figure 4: NCM3722 P1 lysogen gave a higher phage titre as compared to lysates prepared from EMG16 P1 lysogen.** Crude lysates were prepared from NCM3722 and EMG16 harbouring wildtype $P1_{kc}$. Quantification of plaque forming units (in blue) and chloramphenicol-resistant CFU (in orange), representing wildtype P1 phage and P1 transducing unit titres respectively, were carried out on naïve NCM3722 host cells. Crude phage lysates prepared from wildtype P1 lysogenic *E. coli* cell line gave approximately 10 to 15 wildtype P1 bacteriophage per P1 phagemid packaged into transducing unit (Westwater et al., 2002). Each data point is represented a single biological repeat with readings from 4 technical repeats. Data is represented in the form of mean ± SEM.

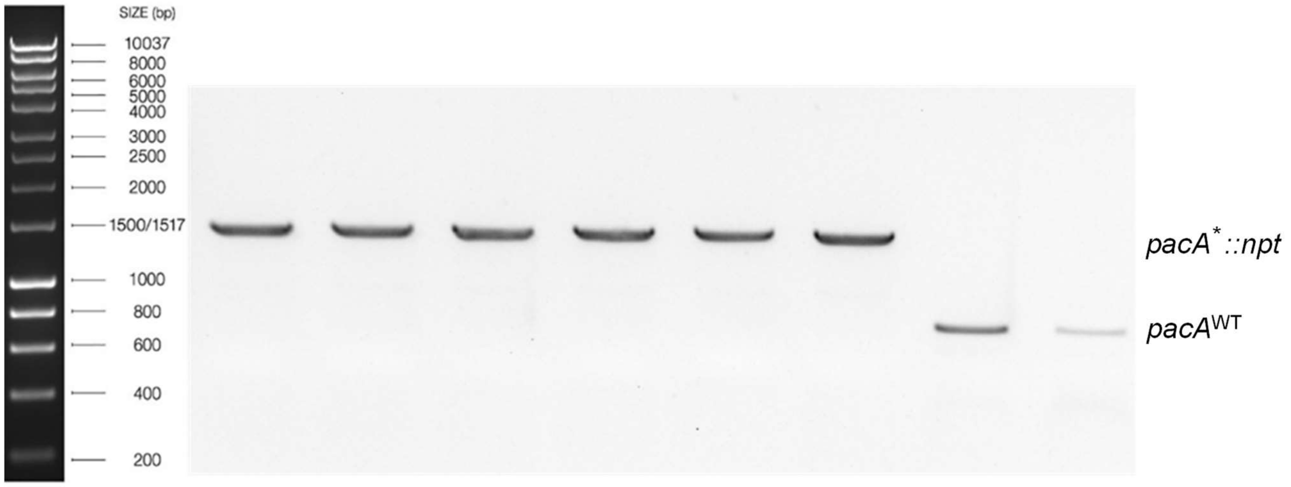


**Supplementary Figure 5: Gel Image showing gel electrophoresis results of the colony PCR reactions, carried out on *pacA*$^{*}$ ::*npt* EMG16 P1 lysogen and wildtype EMG16 P1 lysogen.** *pacA* modified P1 lysogen contains the additional ≈ 850 bps DNA fragment as compared to wildtype P1 lysogen.

**Supplementary Table 1:** Bacterial strains used in this study

| Strain | Description | Remarks/source |
|---|---|---|
| EMG16 *E. coli* K-12 C600 $P1_{kc}$ | *E. coli* P1 lysogen used for phage lysate preparation | CGSC#: 4405 |
| EMG16 *E. coli* K-12 C600 *pacA*$^{-}$::*npt*(kan$^{R}$) $P1_{kc}$ | *E. coli pacA* modified, P1 lysogen with kanamycin resistance cassette retained on the P1 bacteriophage genome | Strain generated in this study |
| *E. coli* NCM3722 | Used for phage lysate preparation, transduction assay and plaque forming units (PFU) quantification | CGSC#: 12355 |
| *E. coli* MC1061::*npt* | *E. coli* that carried a chromosomal copy of the *npt* (neomycin phosphotransferase) antibiotic resistance gene | Strain generated in this study |
| *E. coli* MC1061 | Used as a control to assess the specificity of the chromosomal targeting effect of *cas9*-*npt* phagemid | CGSC#: 6649 |
| *E. coli* TOP10 | Used for routine molecular cloning, assessment of phagemid transduction efficiency | Thermo Fisher Scientific, Waltham, MA, USA |
| *E. coli* BL21 | Used for assessment of phagemid transduction efficiency | Thermo Fisher Scientific, Waltham, MA, USA |
| *E. coli* K-12 MG1655 | Used for assessment of phagemid transduction efficiency | CGSC#: 6300 |
| *S. flexneri* serotype 2a 2457O | Avirulent mutant which contains a transposon insertion in the major-virulence plasmid disrupts *virF* and subsequently prevents activation of the *Ipa* genes essential for bacterial invasion of the host. Used for preliminary assessment of spacer sequence mediated lethality of P1 J72114 phagemids | ATCC: 29903 |
| *S. flexneri* serotype *5a* M90T GFP | An Ampicillin resistant, virulent strain of *S. flexneri* expressing GFP, used for *in vitro* and *in vivo* assessment of spacer sequence mediated lethality of P1 J72114 phagemids | Mostowy *et al.,* 2010[4] |

**Supplementary Table 2** Plasmids and phagemids used in this study

| Plasmid/phagemid | Description | Remarks/source |
|---|---|---|
| BBa_J72114-BBa_J72100 | complete, arabinose inducible phagemid, chloramphenicol resistant, p15a vector, with constitutive *lacZ* expression | Original phagemid construct, a gift from Christopher Anderson (Addgene plasmid #40781)). |
| BBa_J72114.J23115.*gfp* | Complete J72114 phagemid with constitutive promoter Bba_J23115 and RBS Bba_B0030 regulating GFP expression. | Phagemid generated in this study |
| pACYC184-pCas9 | Bacterial expression of Cas9 nuclease, tracrRNA and crRNA guide from *S. pyogenes* | A gift from Luciano Marraffini (Addgene plasmid # 42876) |
| BBa_J72114.*cas9* | Complete J72114 phagemid with constitutive *cas9* expression, tracrRNA and crRNA guide from *S. pyogenes* derived from p*cas9* | Phagemid generated in this study |
| Bba_J72114-pBBR1-c*as9* | Complete J72114.*cas9* phagemid with pBBR1 origin of replication | Used for both *in vitro* and *in vivo* transduction of *S. flexneri* cells |
| pkD46 | Lambda red-mediated recombineering, used for genetic modification of *pacA* gene of P1 bacteriophage | CGSC |
| pSC101.*pacA*$^{*}$::*npt* | *pacA* genetic modification template with kanamycin resistance cassette, pSC101 origin of replication | Plasmid generated in this study |

**Supplementary Tables 3:** DNA sequences of constructs used in this study

| DNA sequence of J72114 phagemid[2] |
|---|
| $Lp_{Sit}$ regulating *pacA* expression[2]<br><br>gaatctggtgtgttaaccgattctacgagtagtcatttgttcattgagtaggaatattgttg |
| *pacA* gene sequence with its native RBS[2]<br><br>cgaaaggaagcataagtgacctgggacgatcacaagaagaattttgctcgcctggcgcgagatggtggttacaccatcgcacagtatgccgccg<br>agtttaatcttaaccctaataccgcacgtcgttatctccgtgccttcaaagaagacaccaggactacggacagccgcaagccaaataagccagtca<br>ggaagccactaaaaagcatgatcattgatcactctaatgatcaacatgcaggtgatcacattgcggctgaaatagcggaaaaacaaagagttaat<br>gccgttgtcagtgccgcagtcgagaatgcgaagcgccaaaataagcgcataaatgatcgttcagatgatcatgacgtgatcacccgcgcccaccg<br>gaccttacgtgatcgcctggaacgcgacaccctggatgatgatggtgaacgctttgagttcgaagttggcgattacctgatagataacgttgaagcgc<br>ggaaggccgcgcgcgctatgttgcgtcggtccggggccgatgttctggaaaccactcttctggaaaagtctctttctcatctccttatgctggagaacg<br>ccagggatacgtgtattcgcctggtgcaggaaatgcgcgatcagcaaaaagacgatgatgaaggtactccgcctgaataccgtatcgcgagcatg<br>ctaaacagctgttccgcgcagataagcagcctgatcaacaccatttacagcatccggaataactatcgaaaagaaagccgggaggcggaaaag<br>cacgctttatctatggggcaagctggcattgttaagctggcatacgaacgaaagcgtgaaaataactggtcagtgctggaagcggctgagttcatcg<br>aggcgcatggaggaaaagtgccgcccctgatgctggagcaaatcaaagccgatctgcgtgctcctaagaccaataccgatgatgaggaaaacc<br>aaacagcatctggcgctccatcacttgaggatctggataaaatcgcgcgagaacgggccgccagccgccgcgctgatgccgcattgtggattgag<br>catcgtagagaagaaattgccgatatcgtcgatacaggtggttatggtgatgtcgatgcggaaggcatatcaaacgaagcatggcttgaacaggat<br>ctggacgaagacgaggaggaagacgaagaagttacccgcaaactgtacggggatgatgattaa |
| Non-coding *repL* sequence (without start codon)[2]<br><br>ttaccctctgaatcctgccggtataccccattgttcgttatctttatttttggctaaaaccgcattaagagcttcgtttaccgtcatgcaatgcggtaggttatc<br>gaagtttgatatcccgccaatatcaggcgaacgcttgttTttcaggtaagcatatttccgcgcagccgcctctactttctgcttgaactcatgtttttgagtg<br>cgtttttggataaccgcagattgtcagcctttgcttttgccttagcgatccatgaagtcaatttttgaggctggttgttccggcaccgccggaaactgatct<br>ttttgtttttttaacttgtgacttcttattctttattgccacgtcatcctgacagggggagggggtatcattttgacatgggggtgtggataaaaaattaaataa<br>agccaatgtcttagcgagaacagctttaaccttggttgccgctgaCgaaatctttaatttgctttctatcagcgcatttttggcttgttgtgcgaaggccaa<br>aaaggatggtgtaaaccggtacaggttagcgcgacgttcacggtgatcgccgataacaatctctacagacagaatacctttgtttacagcttcacgg<br>aatgcacgaacgacggttgattggctataaccagtttctgccgcgatcaggcggtgaggcttgtgaatgaagtattcactggttgttgccgcgagattt<br>gcacattgcgacaggatatgcccggcgctacgggatagaccggagtgtgttacaaagcaggccaattcatagccagaaaaagtaaaatcgcttta |
| Non-coding *cin* sequence (without start codon)[2]<br><br>ccgagttctcttaaaccaaggtttaggattgaaatgatgacgccggaaacttcttataaagcgtggaaacagccacatcatagatgattgcaacctgc<br>ttacgggggatgcccttctccagcaatcgccgcatttgctgccatgtttcttcttggtatttaggccgacgcccacctatacgaccttctgcgcgagctgc<br>atcaagtccagcgcgtgtacgttcaacgataagctcacgttccatttctgccagcgcccccattacgtgaaagaaaaagcgccccattggtgtactg<br>gtgtcgatggagtcagtgagactccggaagttaatgcctctgtcacgcagctcttccaccagcacaactaagtgacgcatgctgcgcccaagacgg<br>tctaacttccatacgaccagggtatcacctctggaaagTtacggagtaccttttttaacccagggcgctcagcctttttgccgctcgccttgtcctcaaa<br>aattagctcacatcctgcgctttcaagagcgtttcgttgtaaagcagtgttttgttcatttgttgatacgcgtacatagcctattag |
| $P_{BAD}$ promoter regulating *coi* expression[2]<br><br>aagaaaccaattgtccatattgcatcagacattgccgtcactgcgtcttttactggctcttctcgctaaccaaaccggtaaccccgcttattaaaagcatt<br>ctgtaacaaagcgggaccaaagccatgacaaaaacgcgtaacaaaagtgtctataatcacggcagaaaagtccacattgattatttgcacggcgt<br>cacactttgctatgccatagcatttttatccataagattagcggatcttacctgacgctttttatcgcaactctctactgtttctccat |
| *araC* with its native promoter and RBS[2]<br><br>ctctgaatggcgggagtatgaaaagtatggctgaagcgcaaaatgatcccctgctgccgggatactcgtttaatgcccatctggtggcgggtttaacg<br>ccgattgaggccaacggttatctcgatttttttatcgaccgaccgctgggaatgaaaggttatattctcaatctcaccattcgcggtcagggggtggtga<br>aaaatcagggacgagaatttgtttgccgaccgggtgatattttgctgttcccgccaggagagattcatcactacggtcgtcatccggaggctcgcgaa<br>tggtatcaccagtgggtttactttcgtccgcgcgcctactggcatgaatggcttaactggccgtcaatatttgccaatacggggttctttcgcccggatga<br>agcgcaccagccgcatttcagcgacctgtttgggcaaatcattaacgccgggcaaggggaagggcgctattcggagctgctggcgataaatctgc<br>ttgagcaattgttactgcggcgcatggaagcgattaacgagtcgctccatccaccgatggataatcgggtacgcgaggcttgtcagtacatcagcga<br>tcacctggcagacagcaattttgatatcgccagcgtcgcacagcatgtttgcttgtcgccgtcgcgtctgtcacatcttttccgccagcagttagggatta<br>gcgtcttaagctggcgcgaggaccaacgtatcagccaggcgaagctgcttttgagcaccacccggatgcctatcgccaccgtcggtcgcaatgttg<br>gttttgacgatcaactctatttctcgcgggtatttaaaaaatgcaccggggccagcccgagcgagttccgtgccggttgtgaagaaaaagtgaatgat<br>gtagccgtcaagttgtcataa |

*coi* with its native RBS[2]

tacagtgaggcataattatggctttcattccaccaaccatcgacgacgttagacattgctctaacgctttatctgtagaccccgccgaaaccgacgctg
cccgcgccattgctgaacactactcaaagatatccaatcaggagtaccgcatcacccaagacgacctggatgatctcactgacacaatcgaatatc
tcatggccactaaccagccagactcacaataa

rrnB T1 terminator after *coi* coding sequence[2]

caaataaaacgaaaggctcagtcgaaagactgggcctttcgttttatctgttgtttgtcggtgaacgctctc

J72114 vector backbone, with p15A *ori*, *cat* gene in blue conferring chloramphenicol resistance[2]

agtcagagtagaatagaagtatcattctcaccaataaaaaacgcccggcggcaaccgagcgttctgaacaaatccagatggagttctgaggtcatt
actggatctatcaacaggagtccaagcgagctcgatatcaaattacgccccgccctgccactcatcgcagtactgttgtaattcattaagcattctgcc
gacatggaagccatcacaaacggcatgatgaacctgaatcgccagcggcatcagcaccttgtcgccttgcgtataatatttgcccatggtgaaaac
gggggcgaagaagttgtccatattggccacgtttaaatcaaaactggtgaaactcacccagggattggctgagacgaaaaacatattctcaataaa
ccctttagggaaataggccaggttttcaccgtaacacgccacatcttgcgaatatatgtgtagaaactgccggaaatcgtcgtggtattcactccaga
gcgatgaaaacgtttcagtttgctcatggaaaacggtgtaacaagggtgaacactatcccatatcaccagctcaccgtctttcattgccatacgaaatt
ccggatgagcattcatcaggcgggcaagaatgtgaataaaggccggataaaacttgtgcttatttttctttacggtctttaaaaaggccgtaatatcca
gctgaacggtctggttataggtacattgagcaactgactgaaatgcctcaaaatgttctttacgatgccattgggatatatcaacggtggtatatccagt
gatttttttctccatttttagcttccttagctcctgaaaatctcgataactcaaaaaatacgcccggtagtgatcttatttcattatggtgaaagttggaacctctt
acgtgcccgatcaaagatccgcaccgccggacatcagcgctagcggagtgtatactggcttactatgttggcactgatgagggtgtcagtgaagtg
cttcatgtggcaggagaaaaaaggctgcaccggtgcgtcagcagaatatgtgatacaggatatattccgcttcctcgctcactgactcgctacgctcg
gtcgttcgactgcggcgagcggaaatggcttacgaacggggcggagatttcctggaagatgccaggaagatacttaacagggaagtgagaggg
ccgcggcaaagccgtttttccataggctccgcccccctgacaagcatcacgaaatctgacgctcaaatcagtggtggcgaaacccgacaggacta
taaagataccaggcgtttccccctggcggctccctcgtgcgctctcctgttcctgcctttcggtttaccggtgtcattccgctgttatggccgcgtttgtctca
ttccacgcctgacactcagttccgggtaggcagttcgctccaagctggactgtatgcacgaaccccccgttcagtccgaccgctgcgccttatccggt
aactatcgtcttgagtccaacccggaaagacatgcaaaagcaccactggcagcagccactggtaattgatttagaggagttagtcttgaagtcatgc
gccggttaaggctaaactgaaaggacaagttttggtgactgcgctcctccaagccagttacctcggttcaaagagttggtagctcagagaaccttcg
aaaaaccgccctgcaaggcggttttttcgttttcagagcaagagattacgcgcagaccaaaacgatctcaagaagatcatcttattaatcagataaa
atatttctaggcctcccctgattctgtggataaccgggatctgtaaggatcaaccactttgtacaagaaagctgggtcgaattgagatccgaacggttt
attacgtacatcaggtaaaactgaccgataagccgctttcttttgggtatagtgtcgtggacagtcattcatctttctgcccctccaaaagtaaaaaccc
gccgaagcgggtttttacgtaaaacaggtgaaactgaccgataagccgctttcttttgggtatagtgtcgtggacagtcattcatctttctgcccctccaa
aagcaaaaacccgccgaagcgggtttttacgtaaaccaggtgaaactgaccgataagccgctttcttttgggtatagcgtcgtggacagtcattcatc
tttccgcccctccaaaagcaaaaacccgccgaagcgggtttttacgtaaatcaggtgaaactgaccgataagccgggttctgtcgtggacagtcatt
catctaggccagcaatcgctcagatcc

DNA sequence of J23110.RBS30.*gfp* construct

J23110

tttacggctagctcagtcctaggtacaatgctagc

RBS30

attaaagaggagaaa

*gfp*

atgcgtaaaggagaagaacttttcactggagttgtcccaattcttgttgaattagatggtgatgttaatgggcacaaattttctgtcagtggagagggtga
aggtgatgcaacatacggaaaacttacccttaaatttatttgcactactggaaaactacctgttccatggccaacacttgtcactactttcggttatggtgt
tcaatgctttgcgagatacccagatcatatgaaacagcatgactttttcaagagtgccatgcccgaaggttatgtacaggaaagaactatatttttcaa
agatgacgggaactacaagacacgtgctgaagtcaagtttgaaggtgatacccttgttaatagaatcgagttaaaaggtattgattttaaagaagatg
gaaacattcttggacacaaattggaatacaactataactcacacaatgtatacatcatggcagacaaacaaaagaatggaatcaaagttaacttca
aaattagacacaacattgaagatggaagcgttcaactagcagaccattatcaacaaaatactccaattggcgatggccctgtccttttaccagacaa
ccattacctgtccacacaatctgccctttcgaaagatcccaacgaaaagagagaccacatggtccttcttgagtttgtaacagctgctgggattacac
atggcatggatgaactatacaaataa

*pacA** modification template, *lpa* coding sequence in blue, flippase recognition motifs (FRT) in red, kanamycin resistance gene (*npt*) underlined with a continuous line, *pacA* coding sequence underlined with a dotted line, *pac* site is in upper case letters, synonymous *pacA** mutations indicated in red lower cases letters

agatccactctctgaatcctgaattctttcgccgtagccgcgctccgcgccggtttgatgaaaaactccagaacgaactctatatgacgcaggacgaa
aaggaacgccgggagcaccagccttgggtaatggcgcgtcaacttttcaataaggtggcccgtcagcaccgtcattacggtaatgccacatccgca
cgtatctgactgtcaaacatgagaattaattccggggatccgtcgacctgcagttcgaagttcctattctctagaaagtataggaacttcagagcgctttt
gaagctcacgctgccgcaagcactcagggcgcaagggctgctaaaggaagcggaacacgtagaaagccagtccgcagaaacggtgctgacc
ccggatgaatgtcagctactgggctatctggacaagggaaaacgcaagcgcaaagagaaagcaggtagcttgcagtgggcttacatggcgatag
ctagactgggcggttttatggacagcaagcgaaccggaattgccagctggggcgccctctggtaaggttgggaagccctgcaaagtaaactggatg
gctttcttgccgccaaggatctgatggcgcaggggatcaagatctgatcaagagacaggatgaggatcgtttcgcatgattgaacaagatggattgc
acgcaggttctccggccgcttgggtggagaggctattcggctatgactgggcacaacagacaatcggctgctctgatgccgccgtgttccggctgtca
gcgcaggggcgcccggttctttttgtcaagaccgacctgtccggtgccctgaatgaactgcaggacgaggcagcgcggctatcgtggctggccacg
acgggcgttccttgcgcagctgtgctcgacgttgtcactgaagcgggaagggactggctgctattgggcgaagtgccggggcaggatctcctgtcatc
tcaccttgctcctgccgagaaagtatccatcatggctgatgcaatgcggcggctgcatacgcttgatccggctacctgcccattcgaccaccaagcga
aacatcgcatcgagcgagcacgtactcggatggaagccggtcttgtcgatcaggatgatctggacgaagagcatcaggggctcgcgccagccga
actgttcgccaggctcaaggcgcgcatgcccgacggcgaggatctcgtcgtgacccatggcgatgcctgcttgccgaatatcatggtggaaaatgg
ccgcttttctggattcatcgactgtggccggctgggtgtggcggaccgctatcaggacatagcgttggctacccgtgatattgctgaagagcttggcggc
gaatgggctgaccgcttcctcgtgctttacggtatcgccgctcccgattcgcagcgcatcgccttctatcgccttcttgacgagttcttctaataactcggta
ccaaattccagaaaagaggcctcccgaaagggggcctttttcgttttggtccggggatcttgaagttcctattccgaagttcctattctctagaaagtat
aggaacttcgaagcagctccagcctacacttgattgcttgcccgttccgggccttttgacatgtgactttcgttaccctcgcgtcaaaaagagtttttacga
aaggaagcataagtgacctgggacgatcacaagaagaattttgctcgcctggcgcgagatggtggttacaccatcgcacagtatgccgccgagttt
aatcttaaccctaataccgcacgtcgttatctccgtgccttcaaagaagacaccaggactacggacagccgcaagccaaataagccagtcaggaa
gccactaaaaagcATGATTATCGACCATTCTAAcGAcCAACATGCAGGcGAcCACATTGCGGCTGAAATtGC
GGAAAAgCAgcGtGTcAATGCCGTTGTCAGTGCCGCAGTCGAGAATGCGAAGCGCCAAAATAAGCGCA
TtAAcGAcCGTTCAGAcGAcCATGACGTAATTACCcgcgcccaccggaccttacgtgatcgcctggaacgcgacaccctggat
gatgatggtgaacgctttgaattcgaagttggcgattacctgatagataacgttgaagcgcggaaggccgcgcgcgctatgttgcgtcggtccggggc
cgatgttctggaaaccactcttctggaaaagtctctttctcatctccttatgctggagaac

| DNA sequence of CRISPR Cas9 construct from pCas9 plasmid (Addgene plasmid #42876) |
|---|
| Promoter sequence of *trans*-activating *crRNA*<br><br>agtattaagtattgttttatggctgataaatttctttgaatttctccttgattatttgttataaaagttataaaataatcttgtt |
| *Trans*-activating crRNA<br><br>ggaaccattcaaaacagcatagcaagttaaaataaggctagtccgttatcaacttgaaaaagtggcaccgagtcggtgc |
| Promoter and RBS regulating *cas9* expression<br><br>atagaatgataacaaaataaactactttttaaaagaattttgtgttataatctatttattattaagtattgggtaatattttttgaagagatattttgaaaaaga<br>aaaattaaagcatattaaactaatttcggaggtcattaaaactattattgaaatcatcaaactcattatggatttaatttaaactttttattttaggaggcaaa<br>a |
| crRNA guide leader sequence in upper case letters, direct repeats in blue and protospacer sequence in black, BsaI sites are underlined.<br><br>TATTTCTTAATAACTAAAAATATGGTATAATACTCTTAATAAATGCAGTAATACAGGGGCTTTTCAAGA<br>CTGAAGTCTAGCTGAGACAAATAGTGCGATTACGAAATTTTTTAGACAAAAATAGTCTACGAGgttttaga<br>gctatgctgttttgaatggtcccaaaactgagaccagtctcggaagctcaaaggtctcgttttagagctatgctgttttgaatggtcccaaaac |

*cas9*, DNA in upper case represent the sequence between *cas9* ORF and the crRNA leader sequence

atggataagaaatactcaataggcttagatatcggcacaaatagcgtcggatgggcggtgatcactgatgaatataaggttccgtctaaaaagttcaa
ggttctgggaaatacagaccgccacagtatcaaaaaaaatcttataggggctcttttatttgacagtggagagacagcggaagcgactcgtctcaaa
cggacagctcgtagaaggtatacacgtcggaagaatcgtatttgttatctacaggagatttttcaaatgagatggcgaaagtagatgatagtttctttcat
cgacttgaagagtcttttttggtggaagaagacaagaagcatgaacgtcatcctatttttggaaatatagtagatgaagttgcttatcatgagaaatatcc
aactatctatcatctgcgaaaaaaattggtagattctactgataaagcggatttgcgcttaatctatttggccttagcgcatatgattaagtttcgtggtcattt
tttgattgagggagatttaaatcctgataatagtgatgtggacaaactatttatccagttggtacaaacctacaatcaattatttgaagaaaaccctattaa
cgcaagtggagtagatgctaaagcgattctttctgcacgattgagtaaatcaagacgattagaaaatctcattgctcagctccccggtgagaagaaaa
atggcttatttgggaatctcattgctttgtcattgggtttgacccctaattttaaatcaaattttgatttggcagaagatgctaaattacagctttcaaaagatac
ttacgatgatgatttagataatttattggcgcaaattggagatcaatatgctgatttgtttttggcagctaagaatttatcagatgctattttactttcagatatcc
taagagtaaatactgaaataactaaggctcccctatcagcttcaatgattaaacgctacgatgaacatcatcaagacttgactcttttaaaagctttagtt
cgacaacaacttccagaaaagtataaagaaatcttttttgatcaatcaaaaaacggatatgcaggttatattgatgggggagctagccaagaagaatt
ttataaatttatcaaaccaattttagaaaaaatggatggtactgaggaattattggtgaaactaaatcgtgaagatttgctgcgcaagcaacggaccttt
gacaacggctctattccccatcaaattcacttgggtgagctgcatgctattttgagaagacaagaagacttttatccatttttaaaagacaatcgtgagaa
gattgaaaaaatcttgacttttcgaattccttattatgttggtccattggcgcgtggcaatagtcgttttgcatggatgactcggaagtctgaagaaacaatt
accccatggaattttgaagaagttgtcgataaaggtgcttcagctcaatcatttattgaacgcatgacaaactttgataaaaatcttccaaatgaaaaag
tactaccaaaacatagtttgctttatgagtattttacggtttataacgaattgacaaaggtcaaatatgttactgaaggaatgcgaaaaccagcatttctttc
aggtgaacagaagaaagccattgttgatttactcttcaaaacaaatcgaaaagtaaccgttaagcaattaaaagaagattatttcaaaaaaatagaa
tgttttgatagtgttgaaatttcaggagttgaagatagatttaatgcttcattaggtacctaccatgatttgctaaaaattattaaagataaagatttttggata
atgaagaaaatgaagatatcttagaggatattgttttaacattgaccttatttgaagatagggagatgattgaggaaagacttaaaacatatgctcacct
ctttgatgataaggtgatgaaacagcttaaacgtcgccgttatactggttggggacgtttgtctcgaaaattgattaatggtattagggataagcaatctg
gcaaaacaatattagatttttgaaatcagatggttttgccaatcgcaattttatgcagctgatccatgatgatagtttgacatttaaagaagacattcaaaa
agcacaagtgtctggacaaggcgatagtttacatgaacatattgcaaatttagctggtagccctgctattaaaaaaggtattttacagactgtaaaagtt
gttgatgaattggtcaaagtaatggggcggcataagccagaaaatatcgttattgaaatggcacgtgaaaatcagacaactcaaaagggccagaa
aaattcgcgagagcgtatgaaacgaatcgaagaaggtatcaaagaattaggaagtcagattcttaaagagcatcctgttgaaaatactcaattgcaa
aatgaaaagctctatctctattatctccaaaatggaagagacatgtatgtggaccaagaattagatattaatcgtttaagtgattatgatgtcgatcacatt
gttccacaaagtttccttaaagacgattcaatagacaataaggtcttaacgcgttctgataaaaatcgtggtaaatcggataacgttccaagtgaagaa
gtagtcaaaaagatgaaaaactattggagacaacttctaaacgccaagttaatcactcaacgtaagtttgataatttaacgaaagctgaacgtggag
gtttgagtgaacttgataaagctggttttatcaaacgccaattggttgaaactcgccaaatcactaagcatgtggcacaaattttggatagtcgcatgaat
actaaatacgatgaaaatgataaacttattcgagaggttaaagtgattaccttaaaatctaaattagtttctgacttccgaaaagatttccaattctataaa
gtacgtgagattaacaattaccatcatgcccatgatgcgtatctaaatgccgtcgttggaactgctttgattaagaaatatccaaaacttgaatcggagttt
gtctatggtgattataaagtttatgatgttcgtaaaatgattgctaagtctgagcaagaaataggcaaagcaaccgcaaaatatttcttttactctaatatca
tgaacttcttcaaaacagaaattacacttgcaaatggagagattcgcaaacgccctctaatcgaaactaatggggaaactggagaaattgtctggga
taaagggcgagattttgccacagtgcgcaaagtattgtccatgccccaagtcaatattgtcaagaaaacagaagtacagacaggcggattctccaa
ggagtcaattttaccaaaaagaaattcggacaagcttattgctcgtaaaaaagactgggatccaaaaaaatatggtggttttgatagtccaacggtag
cttattcagtcctagtggttgctaaggtggaaaaagggaaatcgaagaagttaaaatccgttaaagagttactagggatcacaattatggaaagaagt
tcctttgaaaaaaatccgattgactttttagaagctaaaggatataaggaagttaaaaaagacttaatcattaaactacctaaatatagtctttttgagtta
gaaaacggtcgtaaacggatgctggctagtgccggagaattacaaaaaggaaatgagctggctctgccaagcaaatatgtgaatttttatatttagct
agtcattatgaaaagttgaagggtagtccagaagataacgaacaaaaacaattgtttgtggagcagcataagcattatttagatgagattattgagca
aatcagtgaattttctaagcgtgttattttagcagatgccaatttagataaagttcttagtgcatataacaaacatagagacaaaccaatacgtgaacaa
gcagaaaatattattcatttatttacgttgacgaatcttggagctcccgctgcttttaaatattttgatacaacaattgatcgtaaacgatatacgtctacaaa
agaagttttagatgccactcttatccatcaatccatcactggtctttatgaaacacgcattgatttgagtcagctaggaggtgactgaAGTATATTTT
AGATGAAGAT

**Supplementary Table 4:** Primers for cloning the spacer sequences of *cas9* phagemid crRNAs

| Name of sequence(s) | Sequence (5' to 3')[a] | Remarks |
|---|---|---|
| *npt* forward | AAAC**TTCATCGACTGTGGCCGGCT**G | Chromosomal targeting of *npt* gene of *E. coli* MC1061::*npt* |
| *npt* reverse | AAAAC**AGCCGGCCACAGTCGATGAA** | Chromosomal targeting of *npt* gene of *E. coli* MC1061::*npt* |
| *sigA* forward (G30) | AAAC**ACGACTTTCCCAGTCGGGCT**G | Chromosomal targeting of *sigA* gene of *S. flexneri* (crRNA termed G30) |
| *sigA* reverse (G30) | AAAAC**AGCCCGACTGGGAAAGTCGT** | Chromosomal targeting of *sigA* gene of *S. flexneri* (crRNA termed G30) |
| *pic* forward (G31) | AAAC**GCTTCAGCATTGTTTGAGTC**G | Chromosomal targeting of *pic* gene of *S. flexneri* (crRNA termed G31) |
| *pic* reverse (G31) | AAAAC**GACTCAAACAATGCTGAAGC** | Chromosomal targeting of *pic* gene of *S. flexneri* (crRNA termed G31) |
| *shiD* forward (G34) | AAAC**AATTTCTACTGATTAGATTA**G | Chromosomal targeting of *shiD* gene of *S. flexneri* (crRNA termed G34) |
| *shiD* reverse (G34) | AAAAC**TAATCTAATCAGTAGAAATT** | Chromosomal targeting of *shiD* gene of *S. flexneri* (crRNA termed G34) |
| *shiA* forward (G37) | AAAC**GCATGACTTCTCCGGCTCTC**G | Chromosomal targeting of *shiA* gene of *S. flexneri* (crRNA termed G37) |
| *shiA* reverse (G37) | AAAAC**CGAGAGCCGGAGAAGTCATG** | Chromosomal targeting of *shiA* gene of *S. flexneri* (crRNA termed G37) |

[a]DNA sequences in bold represent the 20 bp spacer sequences.
